# Machine learning analysis of the bleomycin-mouse model reveals the compartmental and temporal inflammatory pulmonary fingerprint

**DOI:** 10.1101/2020.05.22.106690

**Authors:** Natalie Bordag, Valentina Biasin, Diana Schnoegl, Francesco Valzano, Katharina Jandl, Bence M. Nagy, Neha Sharma, Malgorzata Wygrecka, Grazyna Kwapiszewska, Leigh M. Marsh

## Abstract

The bleomycin mouse-model is the extensively used model to study pulmonary fibrosis, however, the inflammatory cell kinetics and their compartmentalisation is still incompletely understood. Here we assembled historical flow cytometry data, totalling 303 samples and 16 inflammatory-cell populations, and applied advanced data modelling and machine learning methods to conclusively detail these kinetics.

Three days post-bleomycin, the inflammatory profile was typified by acute innate inflammation, pronounced neutrophilia, especially of SiglecF^+^ neutrophils, and alveolar macrophage loss. Between 14 and 21 days, rapid-responders were increasingly replaced by T and B cells, and monocyte-derived alveolar macrophages. Multi-colour imaging revealed the spatial-temporal cell distribution and the close association of T cells with deposited collagen.

Unbiased immunophenotyping and data modelling exposed the dynamic shifts in immune-cell composition over the course of bleomycin-triggered lung injury. These results and workflow provides a reference point for future investigations, and can easily be applied in the analysis of other datasets.

## Introduction

Animal models of human disease are an invaluable tool to decipher disease relevant pathomechanisms, to discover therapeutic targets, and to drive translation into clinical practice. To date, the mouse bleomycin-induced lung injury model is the most frequently used animal model to investigate pulmonary fibrosis (B Moore *et al.*, 2013; Della Latta *et al.*, 2015; Tashiro *et al.*, 2017; Biasin *et al.*, 2020). Similar to the human situation, in mice bleomycin exposure is characterized by epithelial damage, inflammatory cell infiltration, and expansion of fibroblasts and myofibroblasts as well as ECM deposition (Biasin *et al.*, 2017, 2020; El Agha *et al.*, 2017; Tashiro *et al.*, 2017; Xie *et al.*, 2018). Although, the bleomycin model does not completely recapitulate human idiopathic pulmonary fibrosis (IPF), it still remains the most common and important animal model to study this disease.

IPF is a severe, rapidly progressing interstitial lung disease with high mortality rates and short median survival of 1.5 - 4 years (Wuyts *et al.*, 2013; Marshall *et al.*, 2018). IPF is characterized by extensive lung tissue scarring, limited inflammation and extracellular matrix remodelling (Meltzer and Noble, 2008). Current treatment options slow the loss of lung function, but are unable to halt or reverse disease progression (Maher and Strek, 2019). Accordingly, there is an urgent unmet clinical need for novel therapies for IPF patients. To date the aetiology and pathogenesis of IPF is still insufficiently understood; however, the role of inflammation remains undeniable yet controversial. The older concept that IPF is an inflammatory driven process has been gradually replaced by the theory of recurrent injury and aberrant repair (Selman, King and Pardo, 2001; Selman and Pardo, 2002; Wuyts *et al.*, 2013). However, multiple inflammatory cells have been implicated in disease pathogenesis, including macrophages (Misharin *et al.*, 2017; Reyfman *et al.*, 2019) and T cells (Todd *et al.*, 2013), which are connected with poorer prognosis (Balestro *et al.*, 2016).

In the bleomycin model, the early phase post bleomycin administration is characterised by acute lung injury and inflammation, which is observed to last between 1-7 days (Peng *et al.*, 2013). This inflammatory phase is followed by active fibrosis, between 7-14 days and late fibrosis between, 21-28 days (Izbicki *et al.*, 2002; Peng *et al.*, 2013; Della Latta *et al.*, 2015; Tashiro *et al.*, 2017). As most studies have only analysed specific cell populations or time points, a comprehensive description of the inflammatory cell kinetics is still missing. For the detection and quantification of inflammatory cells, flow cytometry (FCM) is the method of choice. FCM is able to differentiate and quantify immune cell populations in unprecedented detail, not only from the circulation but also from disease relevant tissue (Misharin *et al.*, 2017; Marsh *et al.*, 2018; Tighe *et al.*, 2019). In contrast to traditional immunofluorescent staining, which generally use 1-3 markers for cell identification, FCM applies multiple markers to simultaneously quantify numerous cell populations at a single cell resolution. Thus, FCM generates large quantities of complex data, where the analysis, visualization and interpretation of which requires sophisticated analysis techniques, such as computational flow cytometry (Saeys, Van Gassen and Lambrecht, 2016; Marsh *et al.*, 2018).

In order to conclusively detail the inflammatory cell kinetics in the bleomycin model, we here assembled historical FCM data from 15 different experiments and applied advanced data modelling, including univariate^[Box 1]^, multivariate^[Box 1]^ and machine learning^[Box 2]^ methods. We show how the combination of advanced data modelling and in-depth immune profiling can detail the dramatic changes in the inflammatory landscape in this model and also serves as a reference point for future studies

### Box 1 Glossary of analysis terms

|  |  |
| --- | --- |
| **robustness** | a measure for how easily outlier values distort results:<br>• average: not robust, a single strong outlier deforms results severely<br>• median: very robust, good results even with almost half of all values being strong outliers |
| **unbalanced** | describes unequal group sizes or missing values, methods assuming balanced groups will have misleading results |
| **positive skew** | asymmetric distribution of data with more small than large values, common in flow cytometry and many other biological measures |
| **data pre-processing** | pre-processing aims to normalise data distribution (i.e. make a bell-shape) by changing all values according to one or several defined mathematical equations<br>• all types of pre-processing can be combined with each other<br>**centering and scaling**<br>cell count differences are not per se reflective of their biological importance, thus centering and scaling reduces the stark differences of cell numbers between the cell populations to allow comparisons of different cell populations. Are vital for multivariate statistical methods, otherwise results will be dominated by cells with highest counts or highest noise<br>• centering: subtraction of a constant from every value, e.g. the mean<br>• scaling: normalize the range of measured values by dividing with a constant e.g. the standard deviation<br>• can be combined, e.g. centering by mean, scaling by standard deviation is z-scaling<br>**transformation**<br>• convert each measured value by a specific, often nonlinear, but defined mathematical function (e.g. $\log_{10}(x)$) to improve distribution<br>• normal distribution is often a prerequisite for specific statistical methods or allows use of more powerful statistical methods (Keene, 1995; van den Berg *et al.*, 2006) |
| **data contaminations** | denotes all kinds of problematic values in the data, such as sample outliers, single value outliers or missing values |
| **outlier** | a value so different from the rest, that it could be for example an analytical error |
| **univariate or multivariate** | univariate methods investigate each measured data on its own (e.g. analysing only CD3<sup>+</sup> T cells irrespective of the 15 other cell populations), while multivariate analyse multiple/all measured data at once (e.g. all 16 cell populations)<br>• univariate methods can dissect in great detail several biological factors (e.g. treatment, substrain) and their interaction, but cannot directly compare different measured data with each other (e.g. is inflammation on a given driven more by T or B cells?)<br>• multivariate methods allow a holistic comparison of various biological factors and their main drivers (e.g. inflammation at day 3 is strongly driven by PMN and less by CD8<sup>+</sup> T cells in BALF Fig 6C), but are limited dissecting several biological factors or their interaction |

### Box 2 Glossary of multivariate methods

|  |  |
| --- | --- |
| **unsupervised or supervised (multivariate)** | unsupervised methods analyse measured data unbiased without any knowledge of biological factors (e.g. treatment, substrain) while supervised methods specifically use biological knowledge<br>• unsupervised methods best detect strongest biological factors, general trends, outliers and comparing impact strength of various biological factors<br>• supervised methods can determine significance of biological factors or in unsupervised methods detected general trends, however the prior knowledge of biological factors necessitates testing and confirmation strategies to avoid overfitting or spurious findings |
| **PCA** | unsupervised method projecting the measured data into new principal components with decreasing variability per component (PC 1 to PC n)<br>• PC1 represents as much variability, i.e. information content (given as percentages), as possible, normally shown in the scores plot on the x-axis<br>• PC2 represents from the remaining variability as much as possible (i.e. PC2 smaller than PC1), shown in the scores plot on the y-axis<br>*scores plot*<br>• each dot represents the whole measured data (i.e. all 16 cell populations) of one sample with closer dots being more similar samples, distant dots are increasingly dissimilar<br>• factors separating groups along PC1 (x-axis horizontal) are stronger, i.e. data is more dissimilar, than separation in PC2 (y-axis vertical) |
| **UMAP** | unsupervised method projecting the measured data (each marker for each cell) into new UMAP components, but optimizing to represent high-dimensional data structure as similar as possible in the low dimensions<br>• interpretation of plots similar to PCA, the nearer sample dots are, the more similar, the further the more dissimilar, often able to handle very large data |
| **machine learning** | algorithms which automatically learn from existing measured data (training data) with the aim to predict unknown data without specific human input<br>• instead of defining a predefined diagnostic cut-off (e.g. AM in BALF below $0.5 \cdot 10^5$ cells is bleomycin treated) a classification model is trained on measured data (e.g. 16 cell counts from BALF of saline and bleomycin treated mice) to predict new, unknown samples (e.g. to verify that the bleomycin challenge and worked)<br>• underlying algorithms can be too complex for human comprehension, even if single elements are simple (e.g. decision tree) due to sheer number of coupled elements (e.g. here 5000 trees in one random forest model) |
| **OPLS-DA** | supervised machine learning method separating predictive variability into the T score (x-axis) from non-predictive into the orthogonal T Score (y-axis) yielding a scores plot similar to PCA<br>• i.e. the T score contains all variability attributed to the given biological grouping (e.g. saline or bleomycin after 3, 14, 21 days), while the orthogonal T score represents technical or inter-individual variability not connected to the biological group<br>• model stability can be assessed by cross-validation and permutation testing, a predictive ability $Q^2 > 50\%$ is considered significant, i.e. high probability to correctly predict new samples |
| **random forest** | supervised machine learning method classifying samples into their given biological group based on measured data using an ensemble of many decision trees<br>• the build-up from many trees hinders simple representations of results, so that several plots are used to assess model stability and main drivers<br>*MDS scores plot*<br>• PCA of the proximity matrix (i.e. similarity of samples) of the random forest model, visually summarizes random forest found similarity/dissimilarity of samples<br>*minimal depth distribution plot*<br>• counts how often the measured data (e.g. CD3<sup>+</sup> T cells) were used in an early node in the trees, sorted according to descending importance<br>• the smaller the mean minimal depth (or the more the distribution contains small values), the more samples were classified based on this measure, the more important this measured data is for a good classification, i.e. drives the difference between groups<br>*rank of mean accuracy decrease in group*<br>• determines importance (e.g. of CD3<sup>+</sup> T cells) by randomly permutating the measured data (i.e. scrambling it nonsensical) and evaluating how the model accuracy decreases, here the mean decrease was ranked between the groups to show which cell type was most important for which group |

## Results

### Pre-processing of flow cytometric data substantially improves statistical analysis performance

Intra-tracheal administration of bleomycin in mice, results in a time-dependent development of fibrosis (Figure 1AB). To comprehensively describe the inflammatory cell kinetics following bleomycin treatment, we assembled and conjointly analysed historical FCM data from 15 independent experiments, which resulted in 159 bronchoalveolar lavage fluid (BALF) and 144 lung tissue samples (Supplementary Table S1). Using standard gating strategies (Misharin *et al.*, 2013; Biasin *et al.*, 2017; Nagaraj *et al.*, 2017; Gungl *et al.*, 2018), a total of 16 cell populations covering the main myeloid and lymphoid cell types (Table 1) were identified (Figure 1C). The aggregation of historical experiments inherently led to an unbalanced^[Box 1]^ experimental design (Supplementary Table S1), which was handled by robust statistical methods^[Box 1]^.

**Figure 1.**
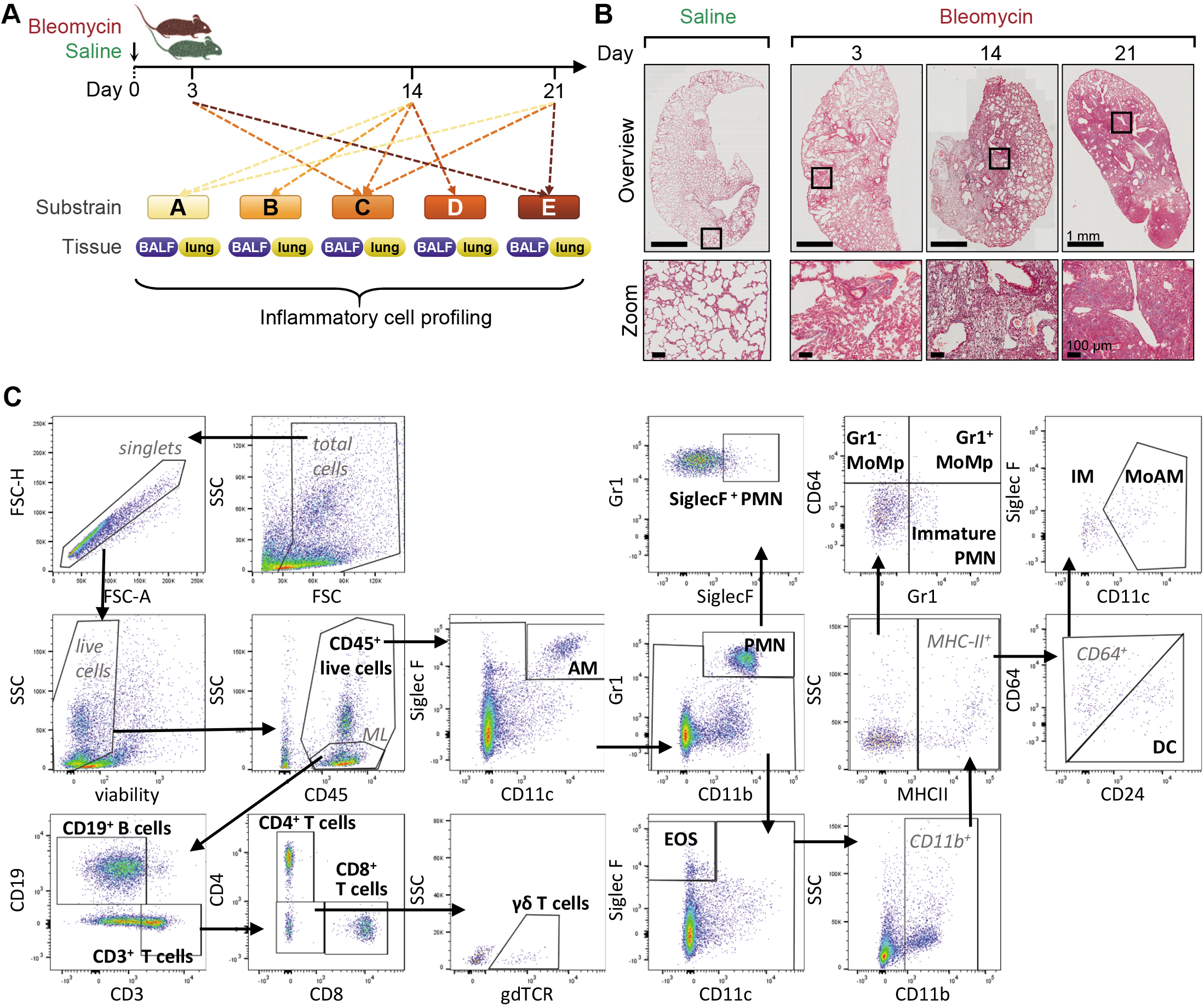
Overview of study design, pathological changes and gating strategy. (A) Historical flow cytometry data from the bleomycin mouse model were pooled and collectively analysed. Samples were collected 3, 14, or 21 days post bleomycin or saline administration from the compartments BALF (159 samples) and lung tissue (144 samples). Five different C57BL/6 substrains were included. (B) Representative Masson’s trichrome staining of lung sections, showing pathologic alterations in the bleomycin model. Zoomed images exemplify the increasing fibrosis accumulation from day 3 to 21 after bleomycin challenge, scale bar represents 1 mm and 100 μm, respectively. High-resolution versions of these images for use with the Virtual Microscope are available as eSlides: VM06176, VM06160, VM06162, and VM06177. (C) Representative flow cytometry gating strategy. The 16 cell populations taken for further analysis are highlighted in bold. Alveolar macrophages (AM), dendritic cells (DC), interstitial macrophages (IM), monocyte-derived AM (MoAM), monocyte-macrophages (MoMp), neutrophils (PMN); forward scatter (FSC), area (A), height (H), side scatter (SSC), monolymph gate (ML). See also Table S1 for overview of group distribution and Table S2 for antibody details.

**Table 1.**
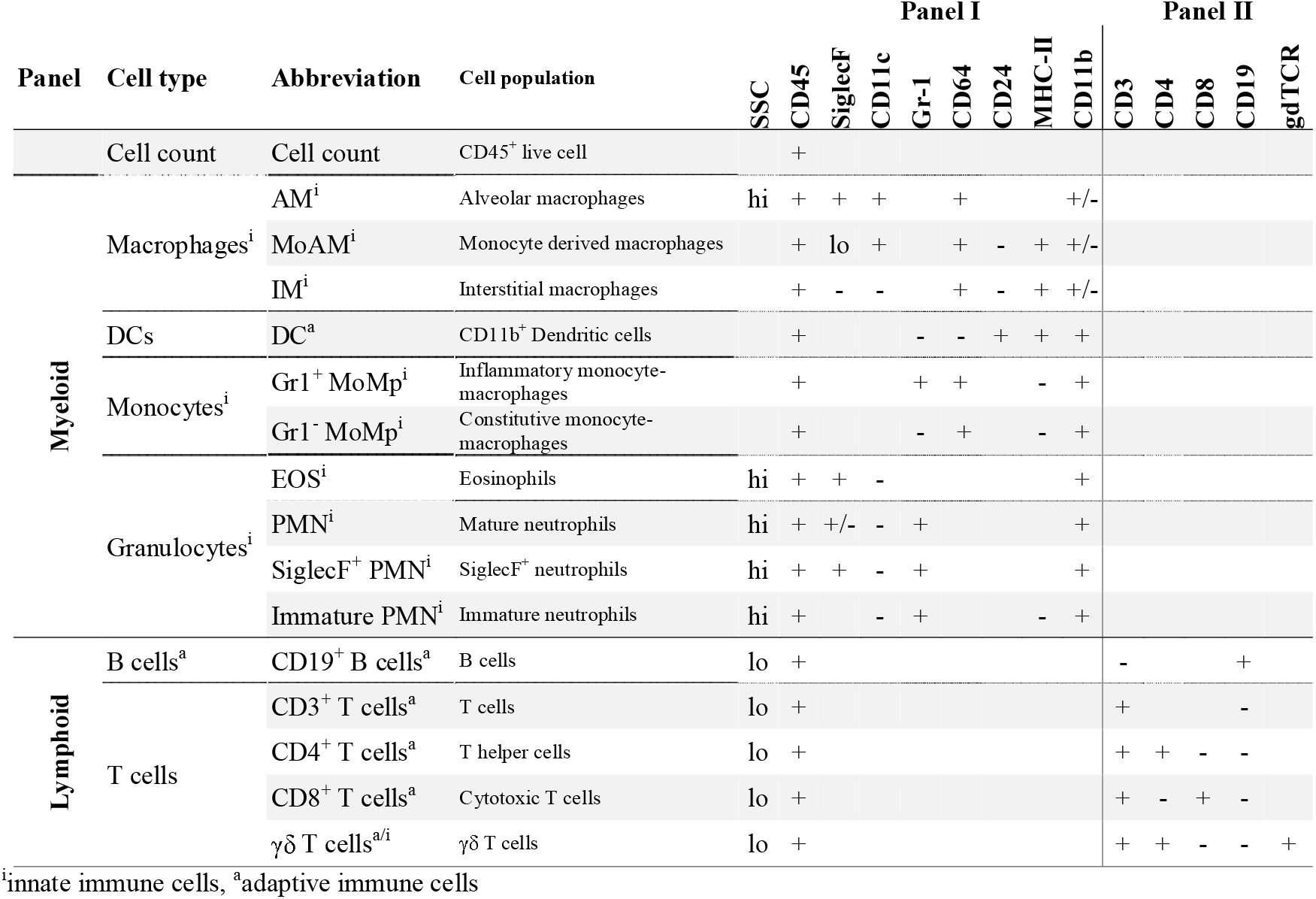
Inflammatory cell identification and corresponding markers.

In both tissues the distribution of all 16 analysed cell populations was significantly non-normal with a positive skew^[Box 1]^ (Fig. 2A, Supplementary Fig. S1 and Supplementary Data 1). To improve distribution we trialled several common transformations; square root, reciprocal, Freeman Tukey, logit, LOG, LOG_x+1_ and 4RT. Only LOG, LOG_x+1_ and 4RT improved data distribution (p_BH_>0.05, Supplementary Data 1). As both LOG and LOG_x+1_ gave virtually equivalent results, but as LOG_x+1_ has additionally the advantage of not introducing missing values for zero value counts, consequent analysis was performed with only LOG_x+1_ and 4RT (Fig. 2AB).

**Figure 2.**
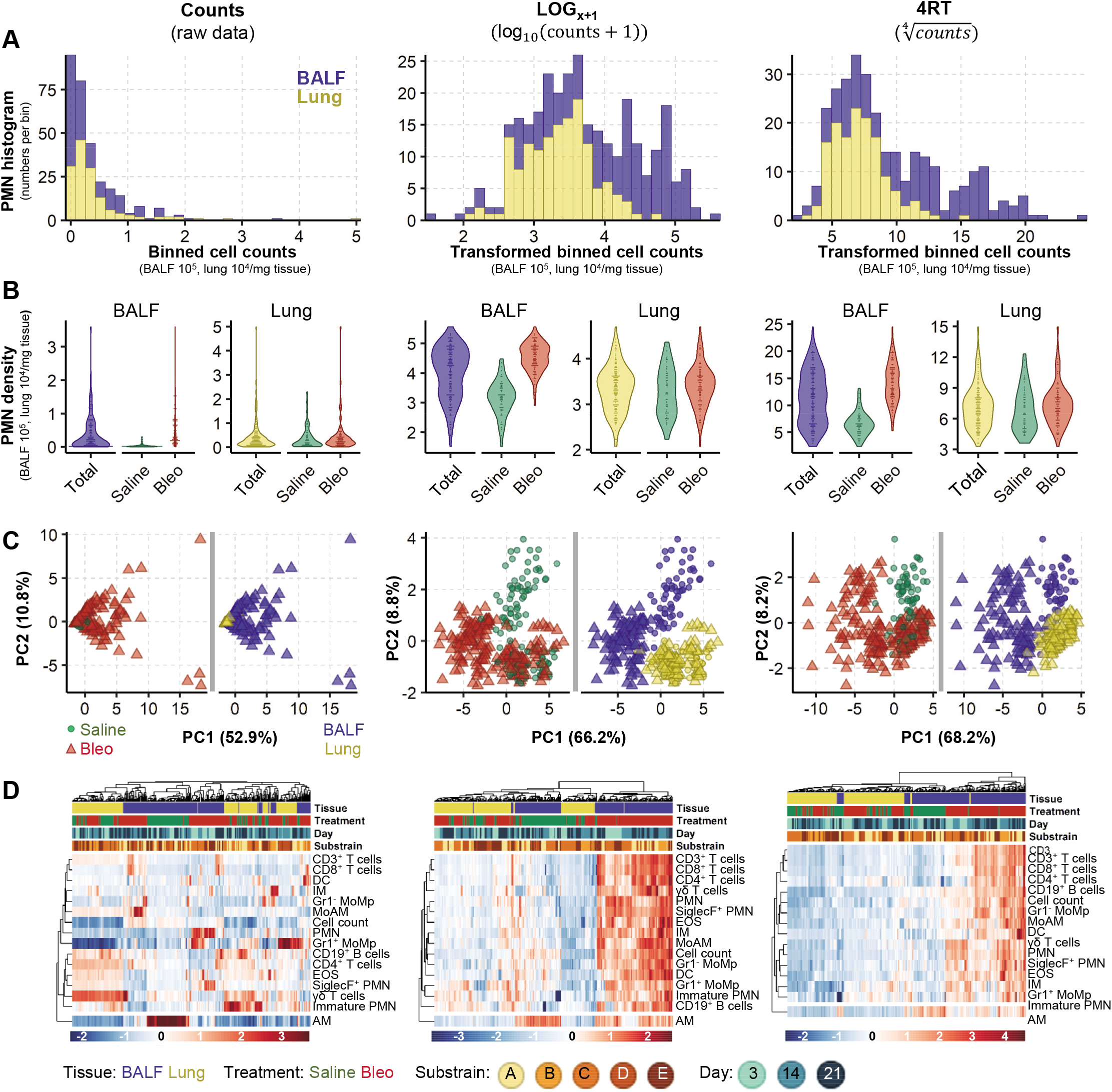
Data transformation improves data distribution and analytical power. Analysis of cell count data (untransformed) or following transformation using LOG_x+1_ or 4RT (fourth root) using 159 BALF and 144 lung samples. Cell counts in BALF are 10^5^ and in lung 10^4^/mg tissue. Examples of data distribution of neutrophils (PMN) as one representative population in BALF and lung samples by (A) Histograms show the frequency of PMN cell counts. Data was grouped into 30 equal intervals (binned cell counts). (B) Violin plots, total represents combined saline and bleomycin samples. (C) PCA scores plots^[Box 2]^ with each point representing the combined inflammatory cell profile (16 populations) in one sample, plots are coloured to highlight different experimental conditions. In B and C, dots represent single sample values. (D) Heatmaps with hierarchical clustering of all 16 analysed cell populations. See also Figure S1 for data distribution and Figure S2 for macroPCA comparison.

### Bleomycin drives strong changes in the inflammatory profile

To identify global changes in the inflammatory cell profile, we first applied unsupervised^[Box 2]^ principal component analysis (PCA^[Box 2]^). This method reduces dimensionality by creating new variables, which successively maximize variance and thereby aids data interpretability. Without data transformation, the scores plot was dominated by single sample differences, which obscured any experimental effects (Fig. 2C, left panel). After transformation, pronounced differences in the inflammatory profile were revealed (Fig. 2C). Both LOG_x+1_ and 4RT substantially improved the performance of the hierarchical clustering, yielding clearer clustering and heatmap results (Fig. 2D). The highest influence on the inflammatory landscape came from the tissue compartment (BALF or lung), causing samples to separate along the first principal component (PC1). The second highest difference was caused by bleomycin, separating samples in the BALF along the second principal component (PC2; Fig. 2C, middle and right panels). Similarly, hierarchical clustering was first driven by the tissue compartment, followed by some weaker subclustering due to bleomycin treatment. The majority of cell populations increased after bleomycin exposure, while alveolar macrophages (AM) decreased (Fig. 2D). We next utilised macroPCA, a robust PCA method able to handle and identify all possible types of data contaminations^[Box 1]^, including strong single value or sample outliers^[Box 1]^ (Hubert, Rousseeuw and Van den Bossche, 2019). MacroPCA results were in good agreement with PCA (Supplementary Fig. S2A), which confirmed that this dataset is free of severe outliers allowing the use of a wide variety of statistical methods (Rousseeuw and Hubert, 2018).

As the strong compartment effect could mask weaker drivers that alter the inflammatory landscape, we analysed BALF and lung samples separately (Fig. 3). In the BALF, bleomycin exposure completely altered the inflammatory landscape, separating samples along PC1 (explaining 63.9 % of the variation in the dataset). However, the bleomycin effect only accounted for 12.4 % of the variation in the lung, separating on PC2 (Supplementary Fig. S2A). Again, macroPCA gave similar results in the analysis of the separate compartments (Supplementary Fig. S2B), reconfirming the absence of critical outliers. Analogous to the PCA findings, hierarchical clustering showed a strong clustering after bleomycin exposure in BALF, which was less clear in lung tissue samples. The influence of day post-treatment and substrain (individual C57BL/6 lines) on cell population changes was less distinct, with only some indication towards a possible sub-clustering due to these factors (Fig. 3B).

**Figure 3.**
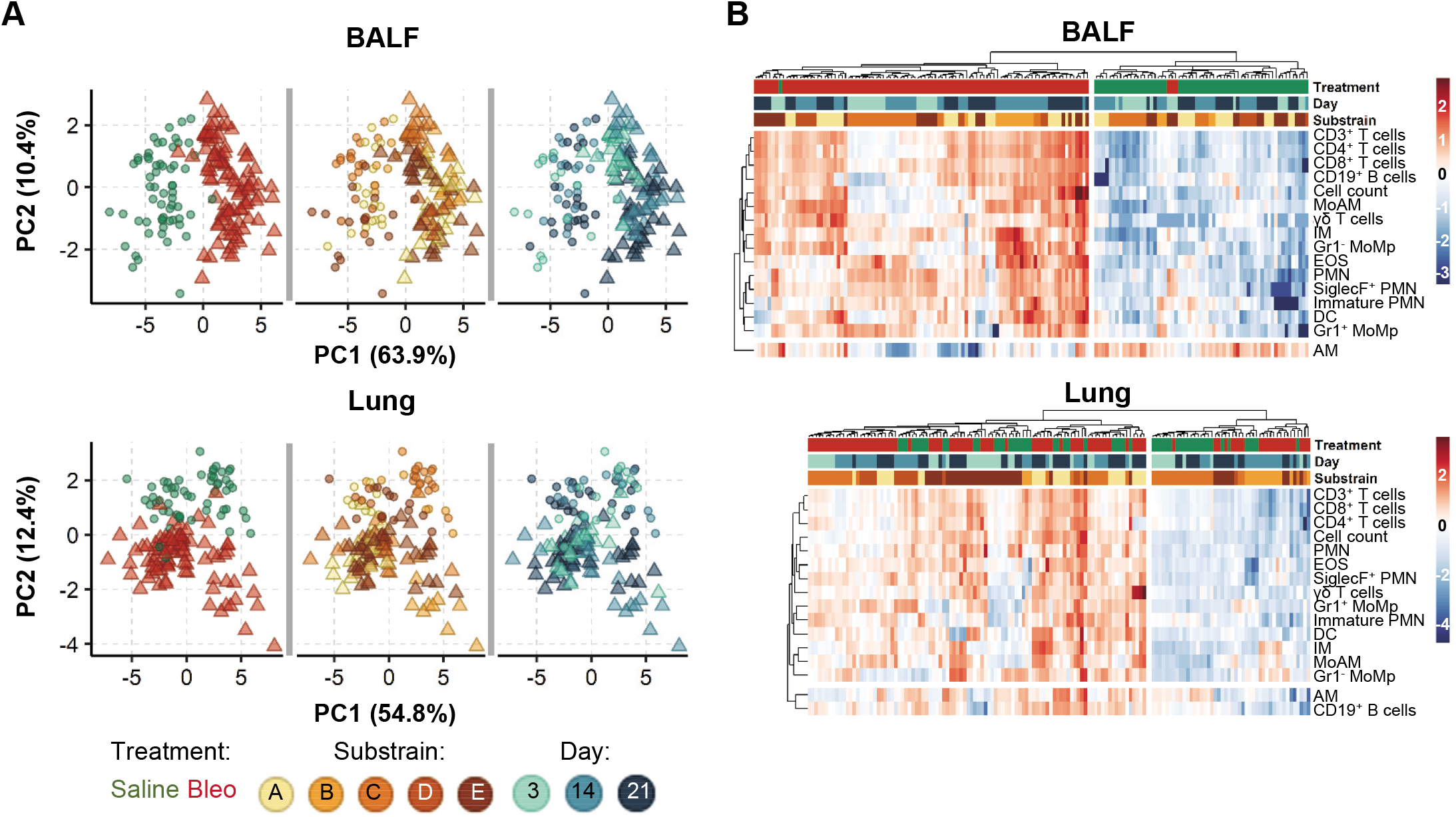
Bleomycin induces stronger changes in the inflammatory profile in the BALF than the lung. The contribution of different biological factors to the inflammatory cell profile as determined by (A) PCA scores plots^[Box 2]^ are coloured to highlight different experimental conditions, and (B) Heatmaps with hierarchical clustering. To aid interpretation heatmaps are split into two main clusters based on dendrogram distances. Colours and shapes represent tissue, treatment (Saline, Bleo), mouse substrain and day post treatment. Cell counts from 16 populations in 159 BALF and 144 lung samples were LOG_x+1_ transformed prior to clustering.

### Modelling of inflammatory cell kinetics with univariate statistical analysis

In order to examine the potential influence of other experimental factors in depth, and to simultaneously control for the unbalanced^[Box 1]^ design arising from the use of historical data, we applied univariate^[Box 1]^ linear mixed models with log_10_-transformation (LOGLME, Supplementary Fig. 3). As the multivariate^[Box 1]^ analysis showed a strong bleomycin effect, the fixed factor^[Box 3]^ *Treatment* {Saline, Bleo} was included in all models^[Box 3]^. Other fixed factors included *Day* {3,14,21} and *Substrain* {A,B,C,D,E}. The addition of each factor, either alone or together and with or without their interaction^[Box 3]^ with *Treatment,* notably improved the fit^[Box 3]^ of all simple models, increasing the goodness of fit and reducing Akaike information criterion (AIC; Supplementary Fig. S3). Thus, both the *Day* post bleomycin exposure and *Substrain* significantly influenced the cellular landscape.

#### Box 3 Glossary of univariate model terms (LOGLME)

|  |  |
| --- | --- |
| **model** | a mathematical equation describing the relationship of measured data to biological factor(s)<br>• you assume that the inflammation, i.e. the CD45<sup>+</sup> cell count, increases with day after bleomycin challenge, then the biological factor is DAY, the measured data is the cell count<br>• a linear model would have the equation: $\text{cell\_count} = a \cdot \text{DAY} + b$ where the fit parameters are a the inclination (steepness of the line) and b the intercept (weight at height=0) |
| **fitting** | finding the parameter values best describing the measured data, often assessed by the residuals |
| **residuals** | difference between fitted value and measured value (in linear models the distance from the measured data dot to the line) |
| **fixed factor** | also called between-subject effect, a biological factor which (possibly) affects the outcome, e.g. treatment or day after treatment |
| **interaction** | the impact of one biological factor depends on the occurrence of another biological factor<br>• e.g. the inflammatory effect of treatment depends on the day after treatment, such as CD45<sup>+</sup> cell count is higher after 14 days than 3 days (in BALF) |
| **random factor** | also called within-subject effect, a factor which (possibly) affects baseline level such as repeated measures from the same source or working in experimental batches<br>• e.g. in one experimental run the cell isolation yielded in all populations higher cell counts than in another experimental run (higher baseline), but does not impact relative findings |
| **simple/mixed** | simple models contain only fixed factors factor, mixed models include random factors |
| **fitted value** | the value suggested by the model from the fitted equation (measured value minus the fitted values is the residual), if the model is correct that would be the real value without measurement error |
| **predicted value** | similar to fitted values the predicted value is suggested by the model equation, but for formerly unknown or not measured points (e.g. CD3<sup>+</sup> T cells day 21 in BALF in supplementary Fig 4B) |
| **overfitting** | the model contains more parameters than the existing data allows to fit well and thus the model will fail to predict new data correctly<br>• e.g. by including irrelevant factors such as mouse colour, tail length, ear size, etc., one could built perfect models without any relevant foundation or prediction of new data |

As each independent experiment could have similarities, the experimental ID was then included as a random factor (~1|Exp_ID). These mixed models significantly outperformed the aforementioned simple models. Finally, complex mixed models (combining the mixed models with the interactions of *Treatment* with *Substrain* or *Day)* notably outperformed all simple models (with or without interactions). The most complex mixed model *[Treatment+Day+Substrain+Treatment:Substrain+Treatment:Day, ~1\Exp_ID]* outperformed all other models, although more prominently in BALF than in lung (Supplementary Fig. S3 A).

As complex models risk overfitting^[Box 3]^, especially in light of the unbalanced design, we then investigated model simplification. We first tested whether it was possible to create one control group of all saline animals. In all mixed and complex models (i.e. with random factor *Exp_ID)* only 4 of the over 10000 investigated pairwise comparisons of a saline subgroup with another saline subgroup had a p_BH_<0.01 in any of the 16 cell types. This means saline treated animals were sufficiently similar to be combined into one control group. Consequently, *Treatment* and *Day* can be then merged into one fixed factor with four groups: Saline (all days) and bleomycin after days 3, 14, and 21, which was termed *SalineDay* {Saline,3,14,21}, generating the simplified model [*SalineDay*+*Substrain*] and the simplified mixed model *[SalineDay+Substrain~1/ Exp_ID].* The performance of the simplified mixed model was slightly lower than in the most complex mixed model, but well within the range of the other top performing mixed models (Supplementary Fig. S3B).

To compare the models in more detail we also directly compared the fitted values^[Box 3]^ of the simplified mixed model with the most complex mixed model. The fitted values from both models strongly correlated (Pearson correlation R^2^>0.96, Supplementary Fig. S3B). This underlines the validity of model simplification and that no unexpected or systematic skew was introduced. As the simplified mixed model *[SalineDay+Substrain~1\Exp_ID]* also gives more easily interpretable results and has a lower risk of overfitting^[Box 3]^, it was chosen to examine the inflammatory cell kinetics underlying bleomycin mouse model.

This model was then applied to explore how individual substrains may influence the kinetics of different inflammatory cells. All mice included in this study are on the C57BL/6 background, however were obtained from different sources e.g. commercial sources (C57BL/6J, substrain A), or are the wild-type littermates from in-house breedings (substrains B-E). Although some lines were inbred for up to 15 generations, all lines produced similar inflammatory responses in both lung compartments, which only differed in magnitude (Supplementary Fig. S4). This consistency allows to read out the compartmental kinetics of each cell population after bleomycin treatment for all substrains combined.

### Inflammatory cell kinetics after bleomycin-induced lung injury are robust and reproducible

Analysis of the inflammatory response in the BALF, identified a non-resolving inflammatory response, with the total number of inflammatory cells continuing to increase over the investigated time course of 21 days. In the lung tissue, inflammation was characterized by an immediate increase at day 3, stagnating at day 14 and mostly resolved 21 days post bleomycin exposure (Fig. 4). This suggests that the inflammatory response is persistent, yet compartment dependent.

**Figure 4.**
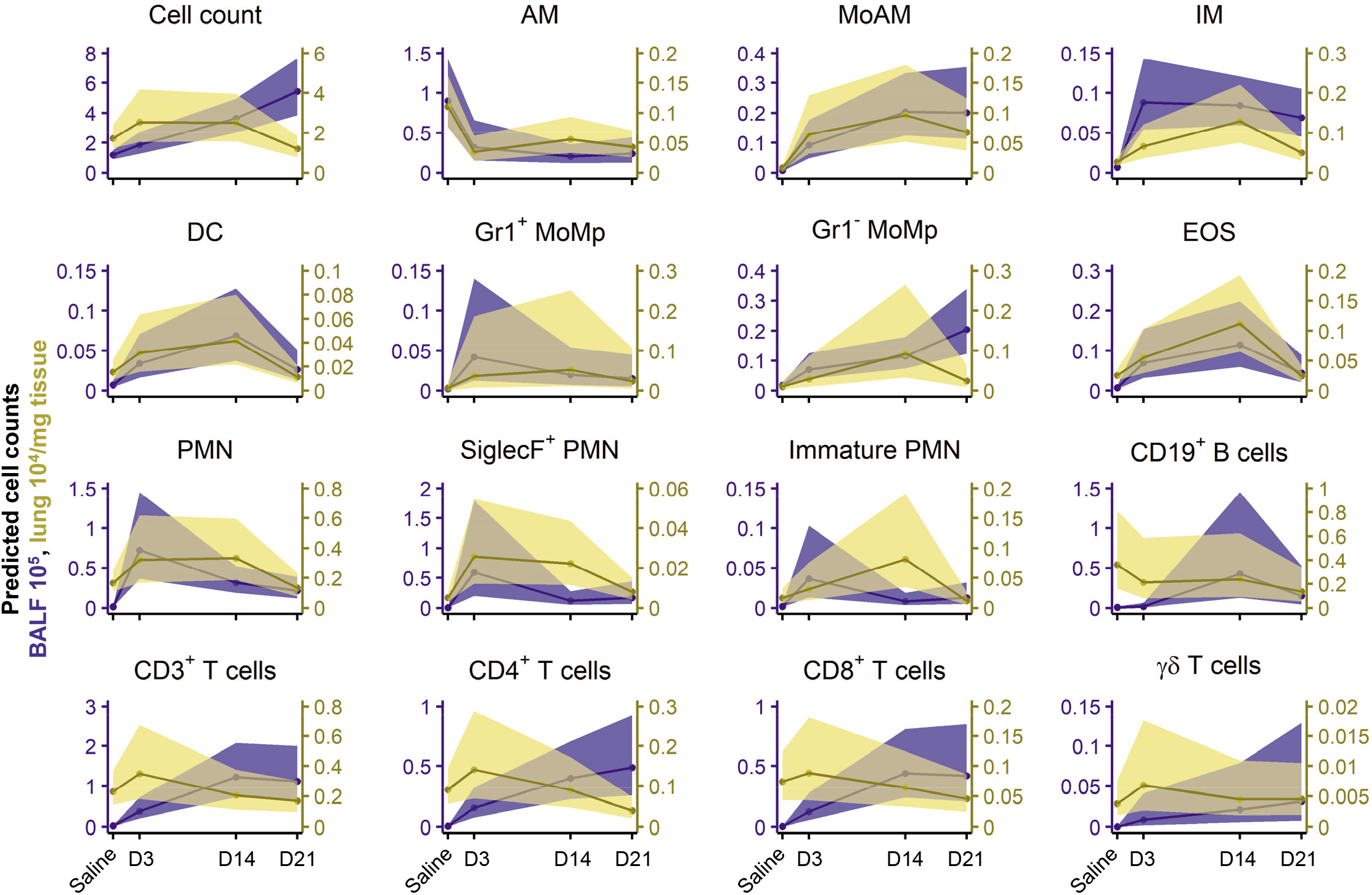
Linear mixed models with log_10_-transformation reveal complex immune cell dynamics occurring in the lung following bleomycin induced lung injury. Plot of back transformed, fitted cell counts (line represents mean ±95 % confidence intervals) using the simplified mixed model *[SalineDay+Substrain~1/Exp_ID]* of LOG_x+1_ transformed cell counts for BALF (counts-10^5^) and lung tissue (counts 10^4^/mg tissue). Animal numbers were in BALF in total n = 159 (Saline 60; 3d 23; 14d 39; 21d 37) and in lung in total n = 144 (Saline 56; 3d 23; 14d 32; 21d 33). See also Figure S3 for model comparisons and Figure S4 for modelling of all 16 cell populations separated by strain.

Early inflammatory changes were mostly dominated by the innate immune system, including both immature and mature neutrophils (immature and mature PMN), monocyte-derived alveolar macrophages (MoAM) and interstitial macrophages (IM). In contrast we observed a concomitant decrease in AM. Interestingly, the (inverted) trajectories of AM were comparable to the rise in MoAM, suggesting a functional replacement by the latter and supports observations in earlier studies (Misharin *et al.*, 2017). Following the rapid increase in the first line responders, PMN, their numbers later stagnated or gradually decreased, and even returned to baseline levels in the lung tissue. We also identified a time-dependent increase in SiglecF^+^ PMN following bleomycin application. These cells have recently been described to be important for cancer progression and murine myocardial infarction (Engblom *et al.*, 2017; Vafadarnejad *et al.*, 2020). Similarly, eosinophils (EOS) and dendritic cells (DC) exhibited a bell-shape response curve. In contrast, monocyte populations (both Gr1^+^ and Gr1^-^ MoMp) exhibited a slower, but consistent, step wise temporal increase, which could be attributed to their contribution to both the innate and adaptive immunity and their role in tissue repair.

At later time points, inflammation was dominated by immune cells from adaptive immunity, with a clear preference to the alveolar compartment. In the BALF, CD3^+^ T lymphocytes (CD4^+^ and CD8^+^ T cells, respectively) had a steep, yet non-resolving, rise early in the inflammatory response. CD19^+^ B cells peaked at 14 days post bleomycin challenge. Interestingly, at the latest investigated time point, 21 days, T cell numbers continued to rise, implicating their involvement at later stages in this model (Fig. 4A).

Taken together, each inflammatory cell population shows dynamic and distinct inflammatory kinetics with some compartmental preference. With time, the involved immune cells shifted from the innate (e.g. PMN) to the adaptive immune system (e.g. T and B cells), with the BALF being more prominently affected than the lung tissue. However, after 21 days the inflammatory profile was still chronically altered in both compartments, suggesting ongoing inflammation.

Based on these results we went back to our FCM data and visualised the kinetics of the most dynamically altered populations using uniform manifold approximation and projection (UMAP)^[Box 2]^ derived plots (Fig. 5A.). UMAP, like PCA, is a dimensionality reduction technique which can utilise the entire flow cytometry dataset (i.e. the positivity or negativity and intensity of each marker for each cell) and reduces this information into a new two-dimensional space. As predicted in our modelling data and easily apparent in the UMAP plots, AM populations strongly decreased following bleomycin exposure, while PMN vastly increased after three days. Adaptive immune cells, such as CD4^+^ T cells, CD8^+^ T cells and CD19^+^ B cells, expanded more at later time points and were virtually absent in saline treated mice (Fig. 5A). Visualisation, using multi-colour immunofluorescence, revealed the presence of CD11c^+^/SiglecF^+^ AM and Ly6G^+^ PMN in saline and day 3 treated mice, while CD4^+^ T cells and CD19^+^ B cells were more prominent at later time-points (days 14 and 21) (Fig. 5B). Interestingly, CD4^+^ T cells were commonly spatially localised to the collagen rich fibrotic lung tissue (Fig. 5B).

**Figure 5.**
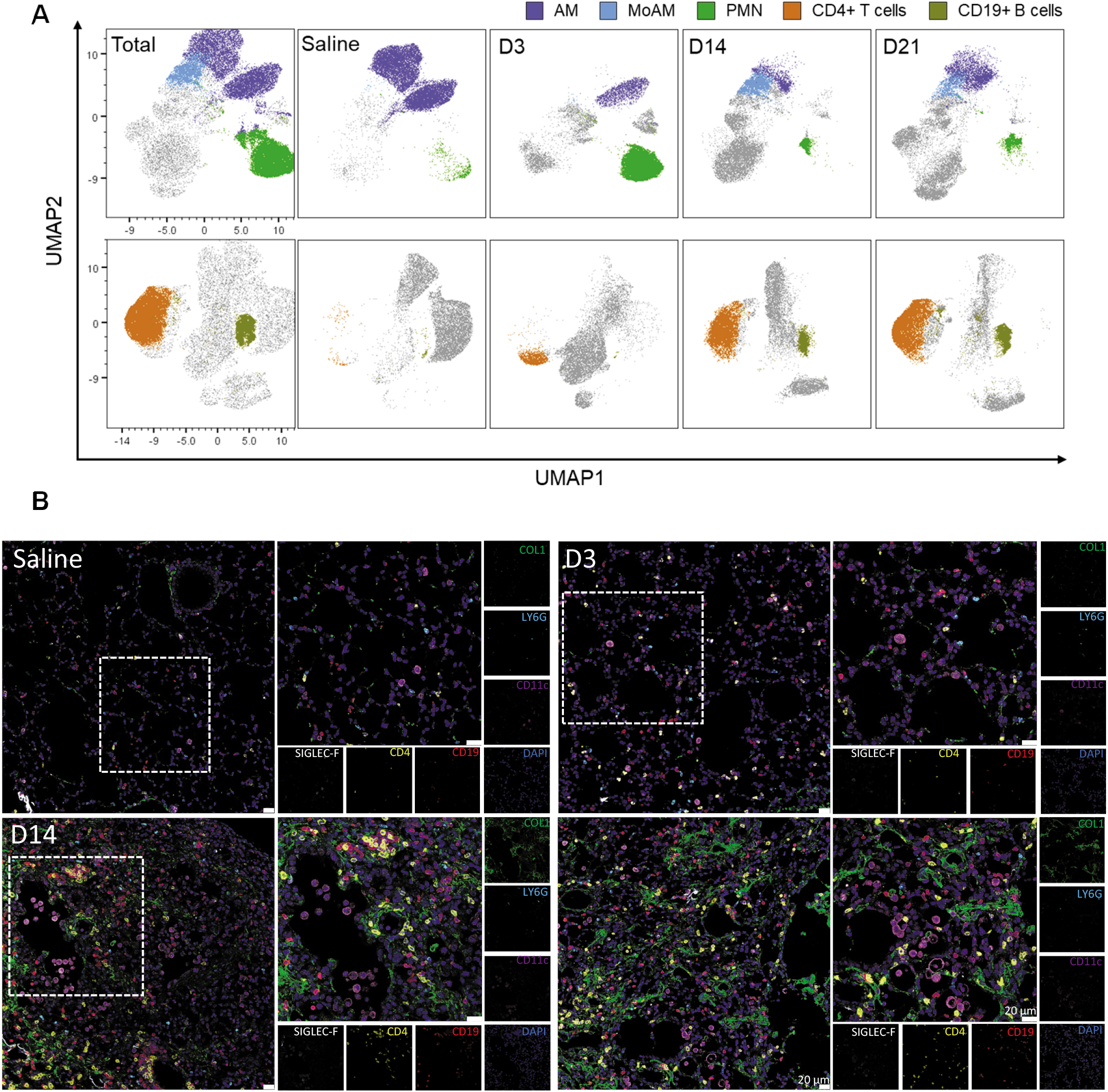
Temporal and spatial localization of inflammatory cell kinetics in BALF and lung tissue. (A) Uniform Manifold Approximation and Projection (UMAP) plots of concatenated CD45^+^ populations (min 3 independent samples with max 10’000 CD45^+^ cells per sample), cells were clustered according to their similarity in signal intensity of all parameters measured by flow cytometry and overlayed with manually gated populations in the BALF. For clarity axis labels are shown only on the first panel of the model. (B) Spatial localisation of alveolar macrophages (AM: CD11c^+^/SiglecF^+^), neutrophils (PMN: LY6G^4^), CD19^+^ B cells and CD4^+^ T cells during the time course of bleomycin challenge. Nuclei are stained with DAPI (dark blue). Representative pictures of three independent mice at each time point. D3, D14 and D21, represent days 3, 14 and 21 postbleomycin treatment, respectively, scale bar represents 20μm. High-resolution versions of these images are available in Virtual Microscope as eSlides: VM06172, VM06173, VM06174, VM06175, respectively. See also Table S3 for antibody details and Table S4 for instrument configuration.

### The inflammatory cell landscape continually evolves following bleomycin exposure

The combination of unsupervised multivariate methods and univariate modelling^[Box 3]^ (LOGLME) identified the kinetics of each cell type with an early innate response followed later by adaptive immune response. However, the question how the entire landscape differs between different timepoints or which cell types define each stage is still open. In order to answer these questions, we applied three robust machine learning^[Box 2]^ approaches.

Our first approach, OPLS-DA^[Box 2]^ separates the dataset into predictive and non-predictive components. Predictive means the ability to discern between groups in the given classification factor, which was here *SalineDay* {Saline,3,14,21}. The OPLS-DA model quality was thoroughly investigated by cross-validation and permutations tests, which showed that in both compartments the models were highly significant (Q2>50 %, p<0.001). Similar to our PCA results (Fig. 3), the inflammatory reaction was more pronounced in the BALF than in the lung, as apparent from a clearer group separation, higher percentages of variability in the predictive component and higher predictive ability (Q2; Fig. 6A). In BALF, the inflammatory landscape at 14 and 21 days post bleomycin were very similar, but very different from the saline controls, while the landscape at 3 days bridged these two poles.

**Figure 6.**
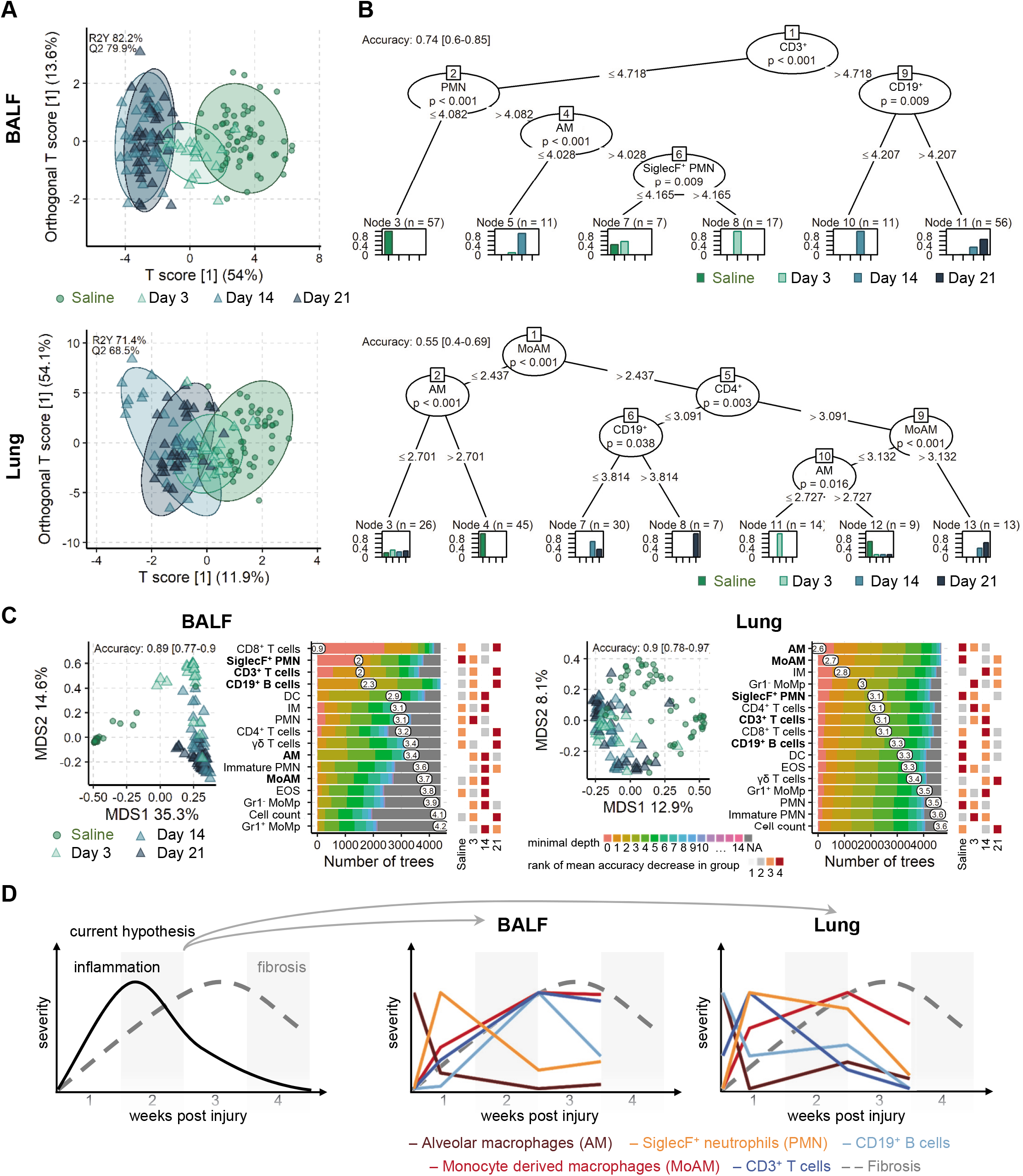
Exploration of inflammatory cell landscape differences with machine learning in BALF and lung tissue. (A) Scores plot of OPLS-DA^[Box 2]^ models per compartment for the factor *SalineDay* {Saline,3,14,21} with 95 % confidence ellipses for each group. The predictive ability of the models Q2 was calculated by 7-fold cross validation and 1000 permutation tests reconfirmed model significance with p<0.001. (B) Conditional inference trees per compartment, showing cell types and cut-offs that define each group; saline, days 3, 14 and 21 post bleomycin treatment *(SalineDay).* Model accuracy was evaluated with a stratified split into 65 % trainings and 35 % test set. (C) Multidimensional scaling (MDS)^[Box 2]^ plot (left panel) of the proximity matrix of random forest^[Box 2]^ models grown with 5000 trees. Model accuracy was evaluated with a stratified split into 65 % trainings and 35 % test set. The distribution of the minimal depth^[Box 2]^ is shown for each cell type according to the number of trees, the mean of the minimal depth is shown (middle panel). The rank of the mean decrease in accuracy^[Box 2]^ within each group is shown for each cell population (right panel). Animal numbers in all models from A-C were in BALF in total n = 159 (Saline 60; 3d 23; 14d 39; 21d 37) and in lung in total n = 144 (Saline 56; 3d 23; 14d 32; 21d 33). Models were based on LOG_x+1_ transformed cell counts for BALF (counts·10^5^) and lung tissue (counts-10^4^/mg tissue). (D) Schematic, abstracted summary of the five lead cell types (highlighted in bold in panel C) and scaled 0 to 1 to highlight relative changes between cell types and compartments BALF and lung tissue.

We next investigated conditional inference trees and random forest^[Box 2]^ models to infer which cell populations were the driving factors behind the group differences. Conditional inference trees in the BALF demonstrated that CD3^+^ T cells levels separated early (Saline, day 3) and later timepoints (days 14 and 21). Separating samples on low and high CD19^+^ B cells distinguishes between days 14 and 21, respectively. On the other hand, low levels of PMN strongly predicts saline treated mice and the combination of low AM and SiglecF^+^ PMN aiding the separation between saline, days 3 and 14 (Fig. 6B). In the lung compartment, both innate cells (MoAM, AM) and adaptive (CD4^+^ T cells and CD19^+^ B cells) were needed to define the different groups. Saline mice were defined by low levels of MoAM and high AM, while bleomycin treatment by high MoAM and CD4^+^ T cells. Similar to the BALF, day 21 was marked by high CD19^+^ levels, while D14 by was defined by lower B cell and MoAM levels (Fig. 6B). A combination of low MoAM and low AM defined day 3.

Random forest models were then used to compare the ability of all cell populations to drive group separation. In agreement with previous results, again group separation was clearer in BALF than in lung, as demonstrated by multi-dimensional scaling plots of the random forest proximity matrix and higher accuracy (Fig. 6C). In BALF especially the adaptive immune cells CD8^+^ and CD3^+^ T cells as wells as the innate SiglecF^+^ PMN differed most, as became apparent from their low minimal depth. Between the different groups high CD8^+^, CD3^+^ and CD19^+^ levels were most predictive for late inflammation while low SiglecF^+^ PMN levels were most predictive for the cellular landscape in saline samples. The random forest suggests some fine but distinct differences between the global inflammatory landscape 14 and 21 days after bleomycin exposure (Fig. 6C). Although both are highly inflamed (OPLS-DA), higher levels of adaptive cells are rather predictive for day 21 (e.g. all T and B cells), while higher levels of some innate cells are more predictive for day 14 than day 21 (e.g. DC, IM, immature PMN, MoAM, EOS) or day 3 (PMN). In contrast, lung models were dominated by macrophage cell populations differing most between the inflammatory stages, foremost the depletion of AM. The random forest models underline that the inflammatory landscape differs notably between lung and BALF.

## Discussion

In this study, we have combined computation FCM, advanced data modelling and machine learning approaches to conclusively define the inflammatory cell kinetics following bleomycin treatment in mice. By combining the data from 15 independent experiments, we amassed very large sample numbers, which were far in excess of those normally found in animal experiments. The aggregation of historical samples inherently led to an unbalanced experimental design, which was handled by sophisticated, robust statistical methods. By using pre-processing techniques such as data transformation, we substantially improved analysis power, which crucially contributed to clearer data interpretation. Changes in the inflammatory profile were dissected using multivariate and univariate statistical methods including linear mixed models with log_10_-transformation. Only by applying these techniques in unison were we able to create the most comprehensive picture of inflammatory cell trajectories to date and characterise the sustained inflammation in the bleomycin model of pulmonary fibrosis. Importantly, these techniques and workflow can be easily applied for analysis of other datasets.

FCM data is normally highly asymmetric i.e. it has many larger values but no values smaller than zero, this non-normal distribution prevents the use of more powerful analysis methods. To re-establish normality we trialled several transformations, but ultimately settled on LOG_x+1_ as it normalised the data distribution, can be easier to interpret and also slightly improved the scedasticity compared to 4RT. Our data modelling approach resulted in a very large sample size, which notably increased statistical power and outweighed the potential drawbacks of added confounding variation from experimental runs or the use of different substrains. Furthermore, when experimental covariance was accounted for as random factor in LOGLME models, the inflammatory profiles in the BALF and lung tissue of all saline treated animals, irrespective of experiment, were sufficiently similar to be combined into one large control group. Secondly, the trajectories of inflammatory cell profiles were found to be consistent for all five substrains, although their magnitudes slightly differed, which is important for experimental reproducibility in light of using different knockout lines or mice sourced from different companies.

The application of unsupervised and supervised as well as multivariate and univariate methods, demonstrated that the majority of cell populations showed consistent trajectories in both compartments. However, the changes for most cells were more prominent in BALF than in lung tissue. This is in part due to fact that in healthy mice the vast majority of cells in the BALF are AM, while in the lung tissue even at baseline conditions, a highly heterogeneous pool of inflammatory cells exists, including macrophages, PMN, T and B cells. The most informative results will be achieved sampling both BALF and lung tissue. The investigation of lung tissue alone could lead to misinterpreting the duration or intensity of inflammation due to weaker changes, while analysis of the BALF alone could potentially miss cell populations that are not normally found within this compartment e.g. interstitial macrophages or to a lesser extent DC. Therefore, deep inflammatory cell phenotyping requires the analysis of both compartments to give the full picture of the inflammatory status of the lung.

Our comprehensive analysis of multiple inflammatory cell population at several time-points, describes the kinetics not only during disease development but also when it is fully established. The initial inflammatory phase after bleomycin exposure was dominated by early responder cell types from the innate immune system of the myeloid lineage. Neutrophils constitute the first line defence of the immune system and consequently show very acute kinetics, being rapidly recruited and also being the first cell type to resolve, visible as pronounced decreases from day 3 to day 14 after the challenge. In contrast, cells from adaptive immune system, such as B and T cells, increased much slower but continue to expand even at 21 days. The worth of subtyping cell populations is apparent by the inverse kinetics displayed within macrophages, which is only possible by using multicolour analysis. We could show that while the numbers of AM quickly decrease, MoAM increased. These contrary trajectories would explain the early observation that macrophages numbers were unchanged in this model (Izbicki *et al.*, 2002), but the closer analysis of macrophage subtypes revealed strong dependent changes, as shown by (Misharin *et al.*, 2013, 2017) and now reconfirmed by our results.

Increasingly macrophage heterogeneity has been suggested to play an important role in the pathogenesis of lung fibrosis and have implications for therapeutic strategies. MoAM undergo marked transcriptional changes during their differentiation in the injured lung tissue. These changes are not only associated with a continuous down-regulation of genes typically expressed in monocytes and up-regulation of genes expressed in AM but also with markedly elevated expression of proinflammatory and profibrotic genes related to M1 and M2 phenotype. This unique transcriptomic signature of MoAM provides an explanation how bleomycin-induced lung fibrosis is attenuated following selective depletion of these cells (Misharin *et al.*, 2017; McCubbrey *et al.*, 2018; Joshi *et al.*, 2020). Interestingly, the existence of common profibrotic pathways in MoAM harvested from mice during fibrosis development and profibrotic macrophages obtained from the lungs of IPF patients has been reported (Misharin *et al.*, 2017; Aran *et al.*, 2019). All these observations strongly suggest that selective targeting profibrotic macrophages, rather than the M1 or M2 phenotype, is more likely to be of benefit in such a complex disease as IPF. The potential contribution of MoAM to the resolution of lung fibrosis remains the subject of future studies, although very recent data supports this hypothesis (Cui *et al.*, 2020). Hence, MoAM could represent a very plastic cell population with distinct functions in different phases of lung fibrogenesis.

Early and late fibrotic stages were characterized by increased numbers of T and B cells in the BALF, while numbers in the lung tissue remained relatively stable, this reflects earlier reports describing the presence of T cells in IPF lungs (Todd *et al.*, 2013; Balestro *et al.*, 2016). Here B cells are of particular interest, as abnormal B cell aggregates have been described in IPF lungs (Marchal-Sommé *et al.*, 2006) and diverse circulating IgG autoantibodies were found in IPF plasma (Ogushi *et al.*, 2001; Kurosu *et al.*, 2008; Taillé *et al.*, 2011). Furthermore, individual auto-immunoglobulins were linked to severity and/or poor prognosis of IPF (Ogushi *et al.*, 2001; Kahloon *et al.*, 2013) thus suggesting the causal role of certain autoantigens in IPF. Accordingly, transcriptome-profiling of lung tissue derived from pirfenidone-treated patients revealed downregulation of B cell related genes (Kwapiszewska *et al.*, 2018). Future studies will, however, demonstrate whether these findings open an exciting new avenue for immunotherapy-based approaches in IPF.

## Recommendations

This study explored fundamental aspects of the bleomycin animal model with good power owing to the high sample numbers so that constructive recommendations can be inferred.

I. In order to ascertain technical success of the experiment we strongly recommend to always include a negative control (saline) and a positive control (bleomycin, transgenic or knock out otherwise untreated) group with each n≥8. Foremost this serves to rate the strength of induced fibrosis and technical quality of the experiment. Statistical power gain is very high for every added sample in the single digit region. An n≥8 leaves some safety margin to stay above the critical level of n=5 to handle the occasional, unavoidable loss of samples due to premature death or technical problems.
II. For more sensitive and pronounced inflammatory readouts the BALF should be routinely sampled together with lung tissue, and both samples subjected to analogous analysis.
III. For statistical analysis we strongly recommend to first investigate distribution and pre-processing (transformation, scaling, centering) paired with unsupervised multivariate methods.
IV. Specific and detailed analysis should be based on the main trends identified in unsupervised models. For detailed investigation of cell specific differences univariate mixed models offer the highest flexibility and insights. Supervised multivariate methods are well capable to confirm wholistic trends and dissect main drivers of these trends. We strongly recommend to pre-processes data before investigating first with unsupervised and second with supervised methods as well as relying on both multivariate and univariate methods, as they complement each other well in their type of generated insights.

## Conclusions

The measurement of inflammatory cellular landscapes in the bleomycin-induced lung-injury mouse model with flow cytometry is very robust and suitable to quantify kinetic changes in multiple cell populations simultaneously. The results allowed to infer recommendations such as to add negative and positive control, apply data pre-processing, combine multivariate and univariate methods and to routinely also investigate BALF. We also found that the unintended development of potential substrains does not per se hinder general reproducibility of results and the approach to adapt bleomycin doses to the current experimental run is viable. This study underlines the relevance of combined analysis for more holistic insights into inflammatory profile changes. Cell populations show quite distinct trajectories in their kinetics. We also conclude that inflammatory cell-based response is active before, during and after manifestation of fibrosis with a shift from the initial innate immune cell dominance towards the adaptive immunity, and importantly inflammatory cell accumulation is not resolved after three weeks.

## Limitations of the study

Despite analysing three independent timepoints, which cover the major stages of the bleomycin model, some timepoints are still missing. However, we consciously wanted to reuse existing experiments and avoid sacrifice of new animals. Future investigation would profit from an expansion, e.g. by inclusion of existing measurements from other groups, to cover also the progression from the initial inflammation towards active fibrosis phase by including analysis between days 3 and 14. Similarly, inflammatory profiling during fibrosis resolution, i.e. after 28 or 35 days, would deliver valuable insights on the involvement of specific subtypes during resolution. From a statistical point of view, the unbalanced study design with differing sample numbers in subgroups is unfavourable, which complicates analysis and loses some power. However, our use of robust methods such as LOGLME and machine learning methods (random forest) were able to overcome these limitations. Although over a dozen independent experimental runs were included, this is not a multi-centric study. Quantitative comparison of results from other laboratories at other sites and other strains/substrains would allow to even better explore bleomycin model system robustness and reproducibility. In this study, manual gating was used to identify different cell populations, thereby including expert knowledge into the analysis and gating specificity was confirmed shown by UMAP overlays. For some populations in the UMAP plots (e.g. AM), the populations were more spread than expected, this was most likely due to do different marker intensity (in this case CD11c) between different experimental runs. The topic of auto-gating is rapidly developing and promises to considerably save hands-on time and foremost the potential to detect rare, otherwise undetected cell subpopulations. The focus of this study was to primarily determine the inflammatory cell kinetics, however to further unravel the role of inflammation and potential therapeutic targets in fibrosis a quantified link of cell subpopulations to fibrotic processes is warranted.

## Supporting information

Supplementary Information, Transparent Methods

## Resource Availability

### Lead Contact

Further information and requests for resources and reagents should be directed to the Lead Contact, Leigh Marsh

### Materials Availability

This study did not generate new unique reagents.

### Data and Code Availability

All data needed to evaluate the conclusions in the paper are present in the paper, the supplementary data is available on Mendeley Data: http://dx.doi.org/10.17632/7j485t986v.1

## Acknowled gements

We are very grateful for the technical assistance from Kerstin Schweighofer, Sabine Halsegger, Thomas Fuchs, Nina Treitler, Eva Grasmann, Camilla Götz, and Betty Flasch, as well as Slaven Crnkovic, Mathias Hochgerner, Chandran Nagaraj and Horst Olschewski for the valuable discussions and advice. Diana Schnögl, Francesco Valzano and Neha Sharma are members of the MolMed graduate programme. Diana Schnögl is funded by FFG, project number 870904 awarded to Leigh Marsh. Francesco Valzano is funded by FFG, project number 874229 awarded to Grazyna Kwapiszewska. Valentina Biasin is funded by Austrian Science Fund (FWF; project number, T1032- B34).

## Author contributions

Conceptualisation, Data curation, Software, Validation, Methodology, N.B. and L.M.M.; Formal analysis and Visualisation N.B., D.S., F.V., and L.M.M.; Investigation, Resources, N.B., V.B., D.S.,

F.V., K.J., B.M.N., N.S. and L.M.M.; Writing – original draft, N.B., V.B., G.K., M.W., L.M.M.; Writing – Review & Editing, all authors; Project administration, Supervision and Funding acquisition,

G.K. and L.M.M.

## Abbreviations

AM: alveolar macrophages
BALF: bronchoalveolar lavage fluid
BH: Benjamini-Hochberg
FCM: flow cytometry
DC: dendritic cells
EOS: eosinophils
IM: interstitial macrophages
IPF: idiopathic pulmonary fibrosis
LOG_x+1_: logarithm to the basis 10 of (x+1)
LOGLME: linear mixed models with log_10_-transformation
MDS: multidimensional scaling
ML: maximum likelihood
MoAM: monocyte derived macrophages
MoMp: monocyte macrophages
OPLS-DA: orthogonal projections to latent structures discriminant analysis
PCA: principal component analysis
PMN: polymorphonuclear neutrophils
UMAP: Uniform Manifold Approximation and Projection
4RT: fourth root

