## Supplementary Information, Transparent Methods for "Machine learning analysis of the bleomycin-mouse model reveals the compartmental and temporal inflammatory pulmonary fingerprint"

### Supplemental Information

#### Supplementary Figures

**A**

| transformation | equation |
|---|---|
| square root | $\sqrt[2]{x}$ |
| reciprocal | $\frac{1}{x}$ |
| Freeman Tukey | $\sqrt[2]{x} + \sqrt[2]{x+1}$ |
| logit | $\ln\left(\frac{x_{map}}{(1-x_{map})}\right)$ with $x_{map} = \frac{(x-Y_{lwr})*((upr-d)-(lwr-d))}{(Y_{upr}-Y_{lwr})} + d$ |
| | BALF $Y_{upr} = 30 \cdot 10^5$, lung $Y_{upr} = 15 \cdot 10^4$, $Y_{lwr} = 0$, $upr = 1$, $lwr = 0$, $d = 0.25$ |
| LOG | $log_{10}(x)$ |
| LOGx+1 | $log_{10}(x+1)$ |
| 4RT | $\sqrt[4]{x}$ |

x – cell counts (BALF $10^5$, lung $10^4$/mg tissue)

**B**

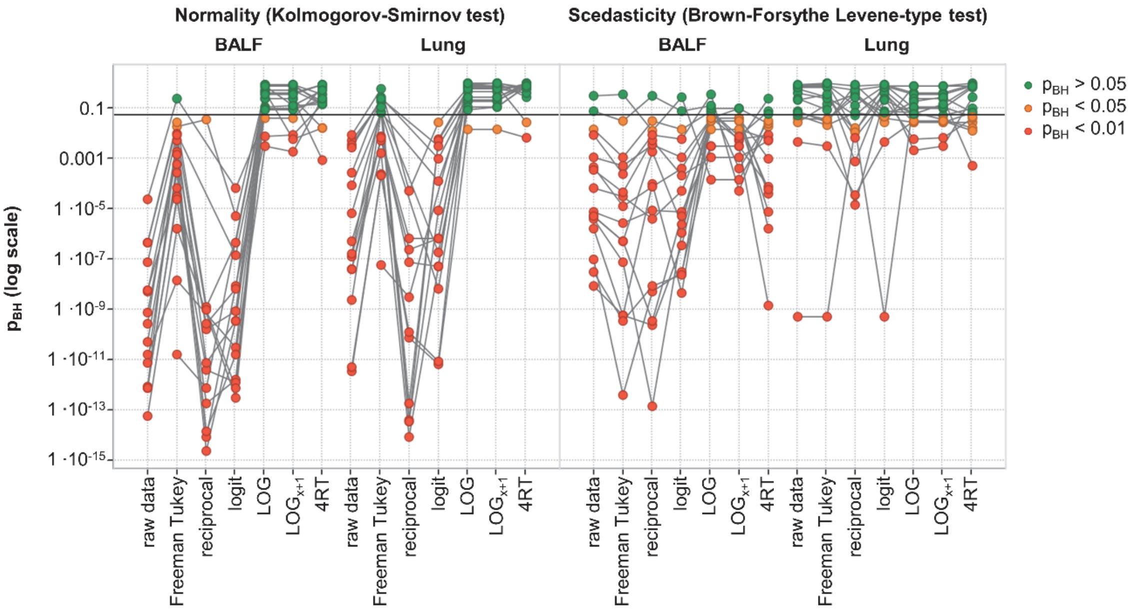


**Figure S1**. Data transformations improve distribution and scedasticity. (A) List of tested data transformations with equations. (B) Normality and scedasticity was tested for each of the 16 cell populations in either 159 BALF or 144 lung samples for each of the transformations. The horizontal line denotes pBH= 0.05. Related to Figure 2.

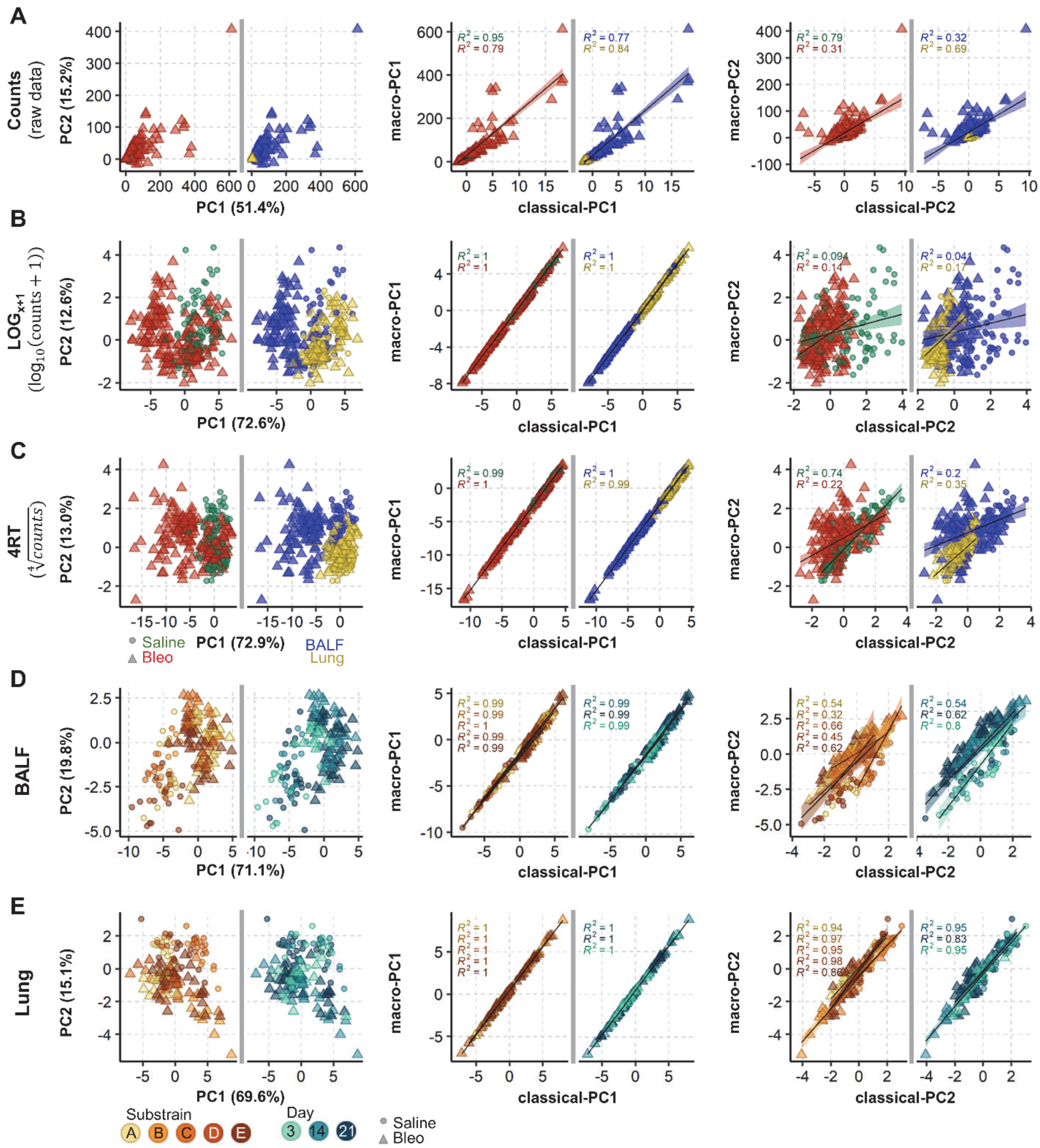

**Figure S2.** MacroPCA and PCA deliver similar results. (A-C) MacroPCA scores plot of combined BALF (159 samples) and lung tissue (144 samples), before (untransformed, (A)) and after data transformation by $LOG_{x+1}$ (B) or 4RT (fourth root; (C)). Samples are coloured to highlight effect of bleomycin (Saline or Bleo) and compartment (BALF or Lung). Middle and right panels show the linear fit of the first two principal components derived from the macroPCA and PCA results. (D-E) Separation of entire $LOG_{x+1}$ transformed dataset into the tissue compartments, BALF (D) and lung (E). Middle and right panels show the linear fit of the first two principal components derived from the macroPCA and PCA results. Samples are coloured to highlight different days and substrains. Shapes are in all plots circles for saline and triangles for bleomycin. Related to Figure 2.

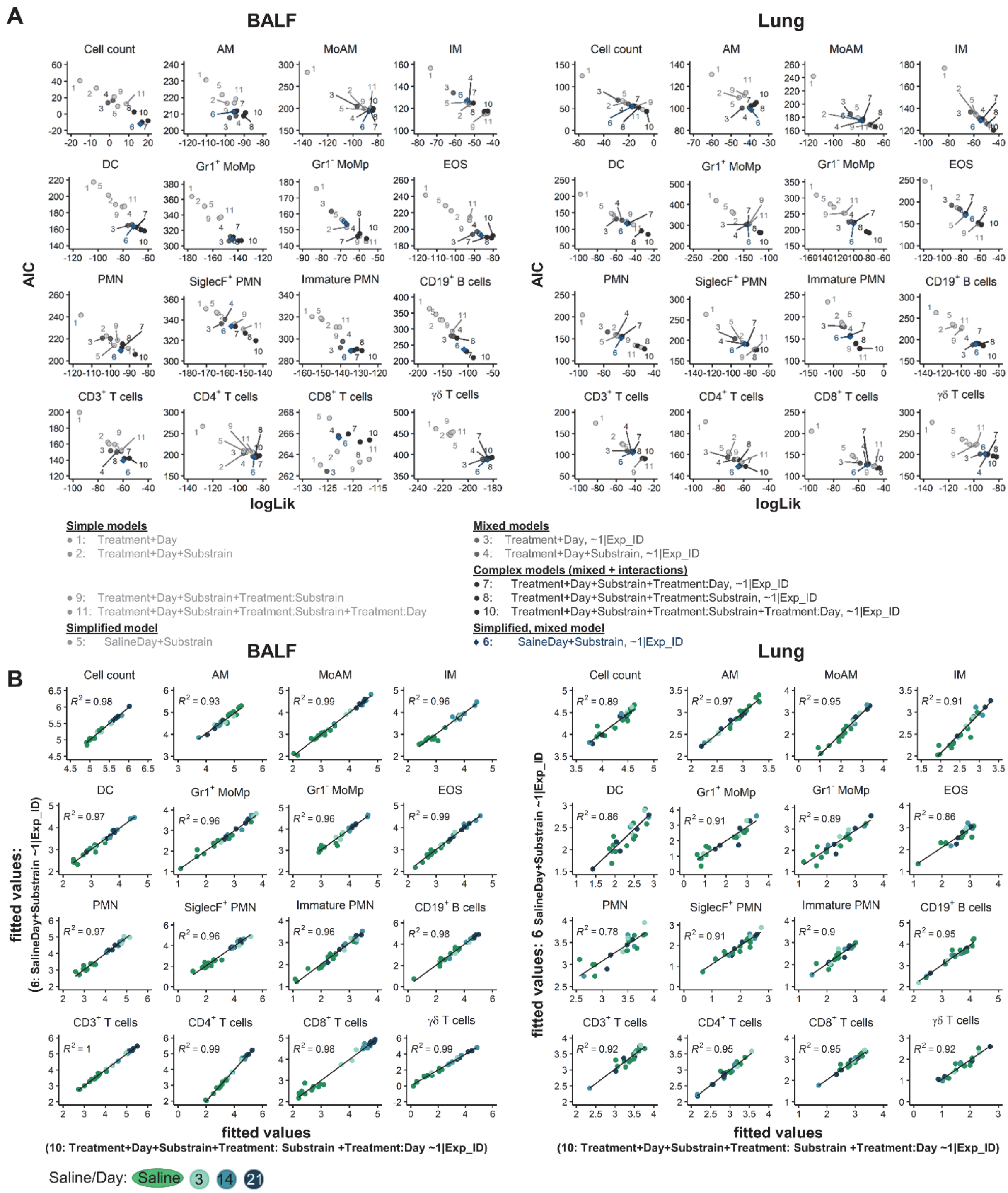


**Figure S3.** Simplified mixed[Box 3] models[Box 3] exhibit best performance. Overview of ANOVA model performances for model selection by: (A) Comparison of model performance by AIC and logLik for all 16 cell populations in BALF and lung, better performance is indicated by lower relative estimate of information loss (AIC; Akaike information criterion) and higher goodness of fit (log-likelihood, logLik). (B) Direct comparison of fitted[Box 3] values (on LOGx+1 scale) of the simplified mixed model versus the most complex mixed model. The Pearson correlation is shown as black line and R2 is given. Related to Figure 4.

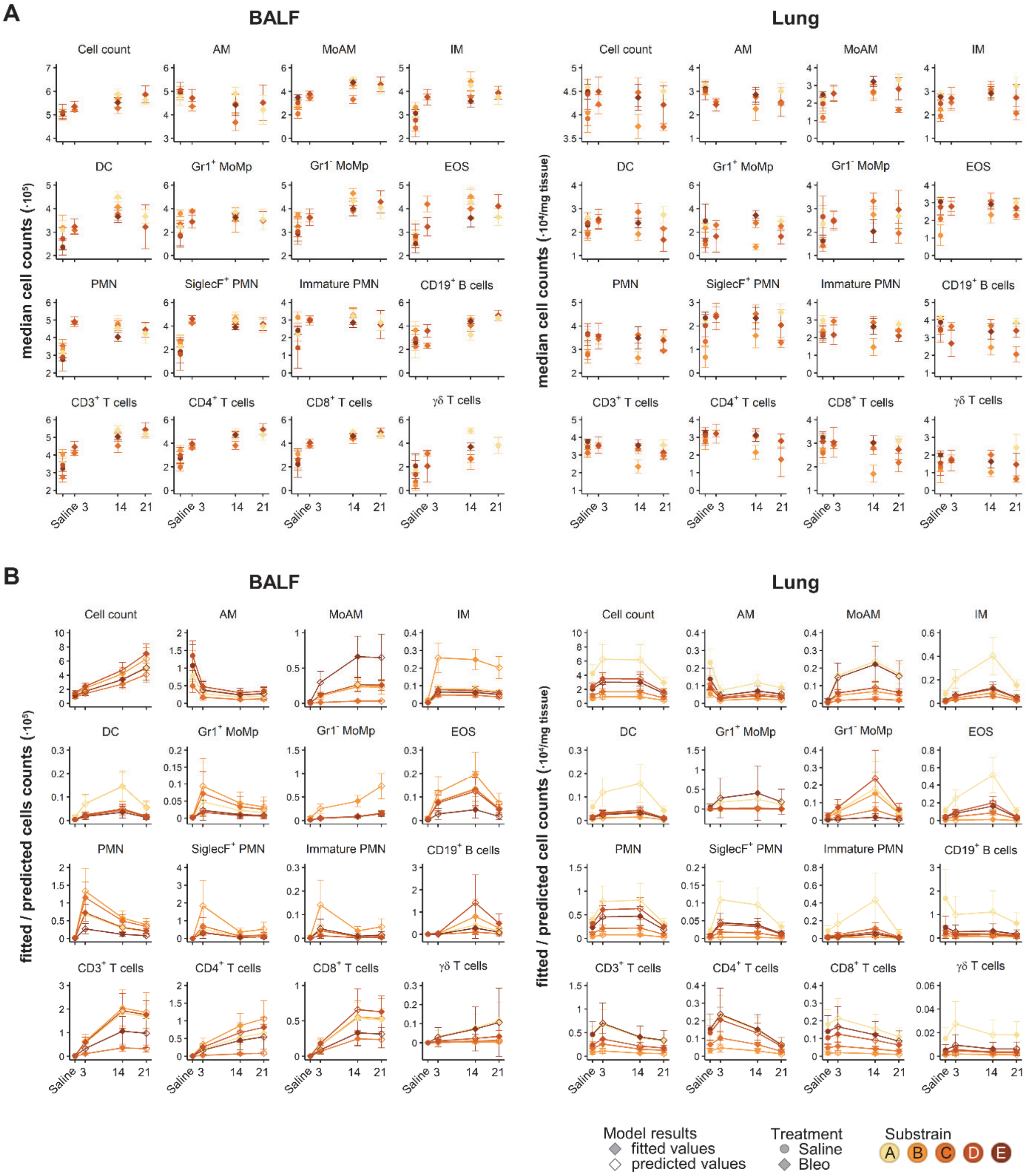


**Figure S4**. Modelling[Box 3] of 16 cell populations in 159 BALF or 144 lung samples reveals complex cell kinetics. Overview of ANOVA model performances for model selection by: (A) Plot of median cell counts at each time point for each substrain and their standard deviation, coloured according to each substrain. (B) Plot of LOG$_{x+1}$ back transformed, fitted[Box 3] or predicted[Box 3] mean cell counts for each substrain and their standard errors from linear mixed models with log$_{10}$-transformation [*SalineDay+Substrain, ~1| Exp_ID*] from cell counts (BALF ·$10^5$, lung ·$10^4$/mg tissue). Related to Figure 4.

**Supplementary Table S1. Overview of group distribution.** Related to Figure 1.

<table>
<tr><th>Substrain</th><th colspan="2">A</th><th colspan="2">B</th><th colspan="2">C</th><th colspan="2">D</th><th colspan="2">E</th></tr>
<tr><th>Compartment</th><th>BALF</th><th>Lung</th><th>BALF</th><th>Lung</th><th>BALF</th><th>Lung</th><th>BALF</th><th>Lung</th><th>BALF</th><th>Lung</th></tr>
<tr><th>Condition</th><th>Saline | Bleo</th><th>Saline | Bleo</th><th>Saline | Bleo</th><th>Saline | Bleo</th><th>Saline | Bleo</th><th>Saline | Bleo</th><th>Saline | Bleo</th><th>Saline | Bleo</th><th>Saline | Bleo</th><th>Saline | Bleo</th></tr>
<tr><td></td><td></td><td></td><td></td><td></td><td></td><td></td><td></td><td></td><td>5|8</td><td>5|7</td></tr>
<tr><td>Day 3</td><td></td><td></td><td></td><td></td><td>8|11</td><td>8|12</td><td></td><td></td><td>3|4</td><td>3|4</td></tr>
<tr><td></td><td></td><td></td><td></td><td></td><td></td><td></td><td></td><td></td><td></td><td></td></tr>
<tr><td></td><td></td><td></td><td>0|4</td><td></td><td></td><td></td><td></td><td></td><td></td><td></td></tr>
<tr><td>Day 14</td><td>4|4</td><td></td><td>0|8</td><td>0|9</td><td>7|13</td><td>4|13</td><td>6|10</td><td>6|10</td><td></td><td></td></tr>
<tr><td></td><td></td><td></td><td>5|0</td><td>7|0</td><td></td><td></td><td></td><td></td><td></td><td></td></tr>
<tr><td></td><td>5|7</td><td></td><td></td><td></td><td></td><td></td><td></td><td></td><td>3|6</td><td>3|6</td></tr>
<tr><td>Day 21</td><td>5|9</td><td>5|9</td><td></td><td></td><td></td><td>6|3</td><td></td><td></td><td>4|8</td><td>4|8</td></tr>
<tr><td></td><td>5|7</td><td>5|7</td><td></td><td></td><td></td><td></td><td></td><td></td><td></td><td></td></tr>
</table>

**Supplementary Table S2. Antibodies, fluorophores and sources for flow cytometry.** Related to Figure 1.

| Panel | Antigen | Label | Company | Catalogue | Clone | Isotype | Identifier | Dilution |
|---|---|---|---|---|---|---|---|---|
| Myeloid | CD45 | FITC | Thermo Fisher | 11-0451-82 | 30-F11 | Rat IgG2b, κ | AB_2753206 | 1:200 |
| | SiglecF | PE | BD Bioscience | 562757 | E50-2440 | Rat IgG2a, κ | AB_2687994 | 1:20 |
| | CD11c | ef450 | Thermo Fisher | 48-0114-82 | N418 | Armenian hamster IgG | AB_1548654 | 1:50 |
| | CD11b | ef506 | Thermo Fisher | 69-0112-82 | M1/70 | Rat IgG2b, κ | AB_2637406 | 1:50 |
| | Gr-1 (Ly6G/Ly6C) | PE-Cy7 | Biolegend | 108402 | RB6-8C5 | Rat IgG2b, κ | AB_313367 | 1:800 |
| | CD64a/b | AF647 | BD Bioscience | 558539 | X54-5/7.1 | Mouse NOD/Lt IgG1, κ | AB_647120 | 1:20 |
| | CD24 | PerCP Cy5.5 | BD Bioscience | 562360 | M1/69 | Rat IgG2b, κ | AB_11151895 | 1:500 |
| | MHC-II | APC-Cy7 | Biolegend | 107628 | M5/114.15.2 | Rat IgG2b, κ | AB_2069377 | 1:400 |
| Lymphoid | CD45 | PerCP Cy5.5 | eBioscience | 45-0451-82 | 30-F11 | Rat IgG2b, κ | AB_1107002 | 1:200 |
| | CD3 | AF700 | Thermo Fisher | 56-0033-82 | eBio500A2 | Syrian hamster / IgG | AB_837094 | 1:50 |
| | CD19 | BB515 | BD Bioscience | 564531 | 1D3 | Rat IgG2a, κ | AB_2738836 | 1:50 |
| | CD8 | PE | Biolegend | 100708 | 53-6.7 | Rat IgG2a, κ | AB_312747 | 1:100 |
| | CD4 | APC | Biolegend | 17-0041-82 | GK1.5 | Rat IgG2b, κ | AB_469320 | 1:100 |
| | gdTCR | ef450 | Thermo Fisher | 48-5711-82 | eBiogL3 | Armenian hamster IgG | AB_2574071 | 1:50 |

**Supplementary Table S3. Antibodies, fluorophores and sources for immunofluorescent staining.** Related to Figure 5.

| Antigen | Host | Brand | Catalogue | Identifier | Dilution |
|---|---|---|---|---|---|
| Collagen I | Goat | Southern Biotech | 1310-01 | AB_2753206 | 1:500 |
| CD4 | Rat | Synaptic Systems | HS-360 017 | AB_2800530 | 1:300 |
| CD11c | Rabbit | Thermo Fisher | PA5-79537 | AB_2746652 | 1:150 |
| SiglecF | Goat | R&D Systems | AF1706 | AB_354943 | 1:500 |
| Ly6G | Rat | Biolegend | 127602 | AB_1089179 | 1:150 |
| CD19 | Rat | eBioscience | 14-0194-82 | AB_2637171 | 1:500 |

**Supplementary Table S4. Instrument configurations.** Related to Figure 5.

| Instrument | Laser lines | Bandpass Filters | | | | | | |
|---|---|---|---|---|---|---|---|---|
| LSRII | 488 nm | 780/60 | 695/40 | 670/14 | 610/20 | 576/26 | 530/30 | 488/10 |
| | 633 nm | 780/60 | 730/45 | 660/20 | | | | |
| | 405 nm | 610/20 | 525/50 | 440/40 | | | | |
| | 355 nm | 530/30 | 440/40 | | | | | |
| Cytoflex S | 488 nm | 690/50 | 525/40 | 488/8 | | | | |
| | 561 nm | 780/60 | 690/50 | 610/20 | 585/42 | | | |
| | 633 nm | 780/60 | 712/25 | 660/20 | | | | |
| | 405 nm | 660/20 | 610/20 | 525/40 | 450/45 | | | |

| Instrument | Parameter | Acquistion seq 1 | Acquistion seq 2 |
|---|---|---|---|
| Leica TCS-SP8 | Pinhole | 67.9 µm | 67.9 µm |
| | PinholeAiry | 1 AU | 1 AU |
| | EmissionWavelength for PinholeAiry Calculation | 580 nm | 580 nm |
| | Excitation Beam Splitter | TD 488/552/638 | TD 488/552/638 |
| Hybrid Detectors | HyD 1 (nm) | | 410 - 460 |
| | HyD 2 (nm) | 492 - 522 | 560 - 571 |
| | HyD 3 (nm) | | 613 - 630 |
| | HyD 4 (nm) | 530 - 548 | 705 - 740 |
| | HyD 5 (nm) | 645 - 675 | |
| Solid state lasers (nm) | 405, Intensity (%): | - | 0.30 |
| | 488, Intensity (%): | 0.30 | - |
| | 552, Intensity (%): | - | 0.40 |
| | 638, Intensity (%): | 0.30 | 0.04 |

#### Transparent methods

##### Animals

All animal experiments were approved by the local authorities (Austrian Ministry of Science, Research and Economics) (BMWF-66.010-0038-II-3b-2013, BMWFW- 66.010/0038-WF/II/3b/2014, BMWFW-66.010/0049-WF/V/3b/2017, 66.010/0177-WF/3b/2017) and were performed in accordance with relevant guidelines and regulations. Wild type groups of 15 independent experiments (unpublished and published (Biasin *et al.*, 2017)) were pooled and analysed. For each experimental run, wild type mice were obtained from Charles River, or bred in-house in case of wild type littermates, and are annotated as separate strains. Overview of all strains and group sizes is given in Supplementary Table S1. All mice were maintained with 12 h light/ dark cycles and had access to water and standard chow *ad libitum*.

##### Bleomycin challenge and animal handling

Male mice (19-32 g body weight, 7-18 weeks old) were anesthetized with isoflurane 2–2.5 % and intra-tracheal administered with bleomycin (Sigma, Vienna, Austria) or saline solution (0.9 % w/v NaCl) using a MicroSprayer® Aerosoliser (Penn-Century Inc., PA, Pennsylvania, USA), as previously described (Biasin *et al.*, 2017, 2020). Each bleomycin lot was titrated to give a comparable response for each strain; dose range was 0.7-3.5 U/kg b.w., Supplementary Data 1). After bleomycin instillation, mice were closely monitored till they completely recovered from anaesthesia. Bleomycin or saline solution administration was performed once and animals were sacrificed after 3, 14 or 21 days.

##### BALF and lung tissue preparation for flow cytometry

Mice were euthanized via exsanguination and the lungs were perfused with phosphate buffered saline (PBS; 137 mM NaCl, 2.7 mM KCl, 10 mM $Na_2HPO_4$, 1.8 mM $KH_2PO_4$, pH 7.2), through the right ventricle. Mice were then lavaged with 1 ml PBS containing the Pierce protease inhibitor cocktail (ThermoFisher Scientific, Vienna, Austria) and 1 mM EDTA. The obtained BALF was centrifuged, washed with 1 ml MACS buffer (2 mM EDTA, 0.5 % BSA in 1X PBS), before being resuspended in 0.5 ml for cell counting and consequent FCM staining. Single cell lung tissue homogenates were performed as previously described (Nagaraj *et al.*, 2017). Briefly, the lower right lobe was weighed, cut into approximately 1 mm pieces and digested with 0.7 mg/ml Collagenase and 30 µg/ml DNAse in RPMI medium supplemented with 10 % FCS, 2 mM glutamine and 1 % penicillin-streptomycin (ThermoFisher Scientific) for 40 min at 37 °C with rotation at 350 rpm. The minced tissue was passed through a 100 µm cell strainer to obtain a single cell suspension. In case of red blood cell contamination, the cell suspension was treated with erythrolysis buffer (2.6 mM $NH_4Cl$, 0.09 M $KCO_3$, 0.6 M EDTA) for 5 min at room temperature. The number of live cells were counted using trypan blue exclusion and then stained with fixable viability stain (ThermoFisher Scientific), washed and then fixed with 1 % paraformaldehyde for 15 min on ice before being resuspended in MACS buffer.

**Flow cytometry**

Single cell suspensions were initially incubated with an Fc-receptor-binding antibody (ThermoFisher Scientific) for 5 min on ice to prevent nonspecific binding. A master-mix containing one of two different antibody combinations against cell surface markers (Supplementary Table S2) was added and cells were incubated for 20 min at 4 °C. For each sample between 30'000 and 300'000 events were recorded on a LSRII Flow Cytometer (BD Biosciences, Vienna, Austria) or Cytoflex S (Beckman Coulter, Vienna, Austria). Samples were analysed either using FACSDiva (BD Biosciences) or FlowJo v10.6.2 (LLC, Ashland, Oregon) software by users blinded to treatment condition. Cells were initially gated on FSC and SSC characteristics and duplexes were removed using FSC-A / FSC-H dot blot, dead cells were gated out using viability exclusion. Cells positive for the pan-leukocyte marker CD45 were taken for further analysis, cell populations were identified using the gating strategy (Fig. 1C and Table 1), as described in the results and based on published studies (Misharin *et al.*, 2013, 2017; Biasin *et al.*, 2017; Gungl *et al.*, 2018; Tighe *et al.*, 2019). A complete description of all antibodies is given in Supplementary Table S2. Cell numbers are reported $10^5$ in the BALF and $10^4$/mg tissue for the lung. Uniform Manifold Approximation and Projection (UMAP) plots were performed in FlowJo, using default settings (nearest neighbours 15, minimum distance value 0.5, Euclidean distance). First, fcs files from at least three individual mice per analysis timepoint were downsampled to max 10'000 events and then concatenated. Manually gated populations were then overlaid on UMAP plots to determine they kinetics.

**Trichrome and immunofluorescence staining**

After BALF, the lungs were inflated with 4 % formalin via the trachea and then paraffin embedded. Slides were cut at 2.5 µm thick and stained with Masson's trichrome according to standard protocols. Slides were scanned and imaged with a Virtual Slides VS120 Microscope and OlyVia Software (both from Olympus, Vienna, Austria). For multi-colour immunofluorescence staining, 2.5 µm paraffin-embedded lung sections were dewaxed and subjected to heat induced antigen retrieval at pH6 (Perkin-Elmer, Waltham, MA) using an antigen retrieval chamber for 15 min at 200 W. Slides were blocked with Perkin-Elmer Antibody Block solution for 20 min in a humidified chamber, and primary antibodies (Supplementary Table S3) were sequentially incubated o/n 4 °C in Perkin-Elmer Antibody Diluent. After washing with TBS-T (274 mM NaCl, 47.6 mM Tris HCl + 2 % v/v Tween20 in $H_2O$) primary antibodies against CD4, SiglecF and CD19 were detected with the Opal Polymer HRP secondary antibody (Perkin-Elmer), using the Opal 540, 620, 690 substrates, respectively. Antibodies against Collagen I, CD11c and Ly6G were used simultaneously and detected with AlexaFluor-conjugated secondary antibodies, donkey anti-goat AlexaFluor488, donkey anti-rabbit AlexaFluor555, chicken anti-rat AlexaFluor647, respectively. Nuclear counterstaining was performed with DAPI solution 1 mg/ml (ThermoFisher Scientific).

**Confocal imaging**

For imaging immunofluorescence stained slides, a Leica TCS-SP8 (DMi8 inverted microscope with a LIAchroic scan head) lightning confocal microscope was used (Leica, Wetzlar, Germany). The acquisition process followed a "sequential workflow" with well-defined settings (shown in Supplementary Table S4). In order to minimize fluorescent overlap the plugin "Channel Dye Separation" of Leica Imaging system was used. The following objectives were used: Plan Fluotar 20x/0.75 multi immersion objective and Plan Fluotar 40x/1.25 glycerol immersion objective. Images were acquired at 2048 x 2048 and a pixel size of 142 x 142nm.

**Statistical analysis**

Data visualisation and statistical analysis were performed with R v3.6.3 (R Core Team, 2020) (using the packages readxl, openxlsx, plyr, stringr, tidyr, reshape, colorspace, RColorBrewer, ggplot2, ggpubr, ggrepel, gridExtra, magrittr, cowplot, plotly, lemon, lawstat, dendsort, pheatmap, cellWise, missMDA, FactoMineR, nlme, emmeans, MetaboAnalystR 2.0, caret, randomForest, randomForestExplainer, partykit, e1071), TIBCO Spotfire v10.9.0, TIBCO, Palo Alto, CA and FlowJo v10 (LLC, Ashland, Oregon). Animals with >30% missing values in the investigated 16 cells populations were excluded from the analysis.

All reported p-values were adjusted for multiple testing according to Benjamini-Hochberg (BH) denoted as $p_{BH}$ (R function *p.adjust*). Distribution and scedasticity were investigated with Kolmogorov-Smirnov test and Brown-Forsythe Levene-type test, respectively ($p_{BH}$ Supplementary Data 1). Seven common transformations were tested: square root, reciprocal, Freeman Tukey, logit (on counts mapped to 0.25-0.75), LOG, $LOG_{x+1}$, 4RT (Supplementary Fig. S1).

Principal component analysis (PCA)[Box 2] analysis (R function *prcomp*) was performed centred and scaled to unit variance (z-scaled) on total cell counts (untransformed, $LOG_{x+1}$ or 4RT transformed). The dataset (303 samples, 16 cell populations) contained no missing values and 1.3 % zeros. MacroPCA analysis (R function *MacroPCA*) was performed centred and scaled to unit variance on total cell counts (untransformed, $LOG_{x+1}$ or 4RT transformed). The number of components was set to cumulatively retain 80 % of explained variance, but to deliver between two and ten components. Hierarchical clustering analysis was performed centred and scaled to unit variance (R function *scale*) on total cell counts, for untransformed data per cell type than samples. $LOG_{x+1}$ or 4RT data was centred and scaled only per cell type. The dendrograms were clustered by Lance-Williams dissimilarity update with complete linkage (R function *dist* and *hclust*) and sorted (R function *dendsort*) at every merging point according to the average distance of subtrees and plotted at the corresponding heat maps (R function *pheatmap*).

Linear mixed models[Box 3] were fitted[Box 3] (R function simple models *gls* or mixed models *lme* with maximum likelihood (ML) using the integrated $\log_{10}$-transformation (LOGLME) reporting back-transformed readouts (R function *emmeans*, option *type = "response"*). This renders the approach non-linear mixed models, however due to the name similarity to the *nlme* function we used LOGLME for clarity. No longitudinal covariance was applied, the mice were sacrificed at each time point. Model selection was based on the forward addition approach and complex models were rechecked by backward dropping of factors. Simple[Box 3] models were constructed using the forward addition approach incorporating the fixed[Box 3] factors *Treatment* {Saline,Bleo}, *Day* {3,14,21} post treatment and the mouse background, *Substrain* {A,B,C,D,E}. The interactions[Box 3], *Treatment:Substrain and Treatment:Day* were include to determine whether the treatment effect depended on the *Substrain* or *Day*. Mixed[Box 3] models additionally included the experimental ID as a random[Box 3] factor (~1|Exp_ID). Complex mixed models were created by combining mixed models with the interactions *Treatment:Substrain* and/or *Treatment:Day*. Models were then simplified by merging all saline samples into one control group generating the simple model [*SalineDay+Substrain*] and by including *Exp_ID* as a random factor the mixed model [*SalineDay+Substrain~1| Exp_ID*]. Due to rank deficiencies arising from the unbalanced[Box 1] design the model *SalineDay:Substrain* was not possible. Criteria for model performance and suitability were lower AIC (Akaike information criterion; relative estimate of information loss), higher log-likelihood (goodness of fit), significance in log likelihood ratio test comparing two models, quality of Q-Q plots and randomness in residual[Box 3] plots (Supplementary Data 1 and Supplementary Fig. S2). Post-hoc pairwise comparisons were readout as back transformed estimates (R function *emmeans*, type = "response") with $p_{BH} \leq 0.05$ being considered statistically significant.

Orthogonal projections to latent structures discriminant analysis (OPLS-DA) [Box 2] on $LOG_{x+1}$ data was performed centred and scaled to unit variance (R function *Normalization* with scaleNorm="AutoNorm" and R function *OPLSR.Anal*) with a standard 7-fold cross validation for the classification factor *SalineDay*. Model stability was additionally verified with 1000 random label permutations.

Conditional inference trees were fit with default settings (R function *ctree*) which limits tree size to include only significant splits avoiding overfitting, so that no further cross-validation or pruning was applied. The random forest[Box 2] (R function *randomForest*) error rates decrease markedly within the first 100 trees and stabilized fully after 1500 to 2500 trees. All reported random forests were grown with 5000 trees to guarantee stability and hyperparameter, mtry (8 in BALF and 2 in lung) was tuned to minimal out-of-bag errors (OOB) (R function *tuneRF*). The model stability and prediction quality (R function *confusionMatrix*) of conditional inference trees and random forest was evaluated by splitting the $LOG_{x+1}$ randomly into trainings/test set (65 % / 35 %) stratified for the classification factor *SalineDay* (R function *createDataPartition*).
